# Uncovering the mesendoderm gene regulatory network through multi-omic data integration

**DOI:** 10.1101/2020.11.01.362053

**Authors:** Camden Jansen, Kitt D. Paraiso, Jeff J. Zhou, Ira L. Blitz, Margaret B. Fish, Rebekah M. Charney, Jin Sun Cho, Yuuri Yasuoka, Norihiro Sudou, Ann Rose Bright, Marcin Wlizla, Gert Jan C. Veenstra, Masanori Taira, Aaron M. Zorn, Ali Mortazavi, Ken W.Y. Cho

**Affiliations:** Department of Developmental and Cell Biology, University of California, Irvine, CA, USA; Center for Complex Biological Systems, University of California, Irvine, CA, USA; Laboratory for Comprehensive Genomic Analysis, RIKEN Center for Integrative Medical Sciences, Yokohama, Japan; Department of Anatomy, Tokyo Women’s Medical University, Tokyo, Japan; Department of Molecular Developmental Biology, Radboud University, Nijmegen, The Netherlands; Division of Developmental Biology, Department of Pediatrics, Cincinnati Children’s Hospital Medical Center, University of Cincinnati College of Medicine, Cincinnati, OH, USA; Department of Biological Sciences, Chuo University, Tokyo, Japan

**Keywords:** Gene regulatory networks, Cis-regulatory modules, ATAC-seq, ChIP-seq, RNA-seq, *Xenopus*, Endoderm, Mesoderm, Linked self-organizing maps, Multi-omic

## Abstract

Mesendodermal specification is one of the earliest events in embryogenesis, where cells first acquire distinct identities. Cell differentiation is a highly regulated process that involves the function of numerous transcription factors (TFs) and signaling molecules, which can be described with gene regulatory networks (GRNs). Cell differentiation GRNs are difficult to build because existing mechanistic methods are low-throughput, and high-throughput methods tend to be non-mechanistic. Additionally, integrating highly dimensional data comprised of more than two data types is challenging. Here, we use linked self-organizing maps to combine ChIP-seq/ATAC-seq with temporal, spatial and perturbation RNA-seq data from *Xenopus tropicalis* mesendoderm development to build a high resolution genome scale mechanistic GRN. We recovered both known and previously unsuspected TF-DNA/TF-TF interactions and validated through reporter assays. Our analysis provides new insights into transcriptional regulation of early cell fate decisions and provides a general approach to building GRNs using highly-dimensional multi-omic data sets.

**Highlights:**

- Built a generally applicable pipeline to creating GRNs using highly-dimensional multi-omic data sets
- Predicted new TF-DNA/TF-TF interactions during mesendoderm development
- Generate the first genome scale GRN for vertebrate mesendoderm and expanded the core mesendodermal developmental network with high fidelity
- Developed a resource to visualize hundreds of RNA-seq and ChIP-seq data using 2D SOM metaclusters.

## Introduction

The cloning of animals by nuclear transplantation and the discovery of transcription factors (TFs) that induce pluripotency, both starting from differentiated cells, are powerful demonstrations that hardwired cell programs can be artificially modified (Gurdon et al., 1958; Takahashi and Yamanaka, 2006). Understanding the transcriptional control of cellular differentiation programs, and their plasticity, is fundamentally important in biology and regenerative medicine. In recent years, there has been a greater realization that simple linear pathways of transcriptional regulation, of genes acting through their downstream products is insufficient to explain complex biological phenomena. This is because genes function in complex and hierarchical networks and the emergent properties of these networks ultimately generate biological outcomes (Levine and Davidson, 2005; Davidson, 2010). These gene regulatory networks (GRNs) are visualized as a network diagram by displaying regulatory interactions as lines interconnecting transcription factors that control gene expression. Thus, identifying network structure, function, and activity is a necessary step toward comprehending the causes of cellular states and behaviors, during both embryogenesis and in adults, as well as understanding phenotypes in disease.

Around 15 years ago, efforts were made to compile the available molecular data into GRNs describing mesendoderm and Spemann organizer development in *Xenopus* (Loose and Patient, 2004; Koide et al., 2005). Arguably, these developmental events represent the earliest and most influential developmental periods in metazoan organisms, leading to large-scale morphogenetic changes and body axis formation, as well as the establishment of the organ and tissue primordia that form the complex adult organism. Most recently, a highly-curated interactome map was generated based on over 200 publications, (Charney et al., 2017a), which represents the most thoroughly examined mesendoderm GRN in any chordate. The *Xenopus* mesendoderm GRN revealed that germ layer specification and subsequent events are not programmed by molecules acting in a linear fashion, but instead are controlled by a set of TFs acting in a complex network. Additionally, previous work has suggested that critical aspects of the mesendoderm GRN are conserved in all vertebrates (Zorn and Wells, 2009). Therefore, a highly robust *Xenopus* GRN is likely to inform conserved paradigms in human development and strategies to direct human cell differentiation.

While the current mesendoderm GRN has revealed critical principles governing early embryonic development including feedback, feedforward, and mutual exclusion mechanisms, a particularly notable finding is the prominent presence of the coherent feedforward loop (Charney et al., 2017a; Paraiso et al., 2020), which suggests that temporal regulation of the cascade of gene expression is extremely important to the process. Despite these advances, the mesendoderm GRN is still far from complete. This is in part because, in the past, network connections were not fully embedded within the larger regulatory architecture nor included temporal and spatial data. Therefore, the GRN provided limited previews of the selected interactions that were chosen with *a priori* knowledge of a given gene behavior. Thus, the mesendoderm GRN is likely to miss many important potential interactions and the involvement of TFs. An alternative approach to overcome this limitation is to generate a GRN based on a combination of computational methods with extensive perturbation analysis. Availability of large-scale functional genomics datasets in recent years allows us to test the utility of such an approach. However, one difficulty is the potential to produce numerous putative interactions that may contain many false positives or non-functional interactions. This concern is supported by ChIP-seq analyses, which often uncover tens of thousands of TF-bound sites, but only a fraction of such sites directly affect gene expression (so-called “neutral binding”) (Li et al., 2008; Kvon et al., 2012). Therefore, it would be extremely valuable if one can develop an approach that embraces the scale of the genomic data, while minimizing false positive interactions. Though other methods have been developed to solve this problem through refinement of peak calling of ChIP-seq datasets (Bardet et al., 2013), we hypothesized that integration of genetic perturbation data types into the analysis of chromatin datasets would allow for a more informed identification of functional binding sites, and by extension gene regulatory network connections.

Given the current accumulation of large genomic data sets, computational GRN inference has become a fairly popular field of research. One common way that these methods operate is using co-expression matrices to build “influential” GRNs (reviewed in Delgado and Gomez-Vela, 2019), which rely on correlations between genes rather than direct mechanistic regulation between TFs and target genes. The vast majority of GRN inference studies using high-throughput data build networks of this type. For the current work, we wished to build a “mechanistic” GRN, so we sought to only find direct connections that engage *cis*-regulatory regions. This is extremely difficult in practice using only one type of data (reviewed in Hu et al., 2020). Some recent predictive algorithms use multi-omic data to build lists of putative functional enhancers (Sethi et al., 2020; Xiang et al., 2020), but they do not incorporate TF binding data to determine whether the TF can directly regulate these enhancers. Others focus on integrating single-cell data types, such as Seurat/Cicero (Pliner et al., 2018; Stuart et al., 2019), which was designed specifically to convert scATAC-seq data into scRNA-seq-like data, with regular scRNA-seq data. Even after applying this approach, it is only able to create “influential” GRNs from gene correlation matrices. Another method, LIGER (Linked Inference of Genomic Experimental Relationships) (Welch et al., 2019) also converts scATAC-seq data into scRNA-seq-like data to smooth the scATAC-seq data to perform cell clusterings. These approaches illustrate the challenges of gathering and analyzing highly-dimensional chromatin accessibility data sets. Here we describe the use of widely available bulk data rather than single-cell data for constructing mechanistic GRNs.

We adapted our linked self-organizing map (SOM) method (Jansen et al., 2019) to the multiple data types available for mesendoderm development: ChIP-seq, ATAC-seq, and RNA-seq of wild-type (temporal and spatial) and perturbation conditions. SOMs (reviewed in Kohonen, 2001) are a type of unsupervised neural network, which train on a set of data to generate a lowdimensional representation, or a 2D “map”. Previous works have successfully used the SOM’s remarkable ability to generate robust clusters (Kiang and Kumar, 2001) by incorporating them into the analysis of highly-dimensional genomic data. For example, SOMs have identified complex relationships between genes and genomic regions in multiple cell types in human and mouse (The ENCODE Project Consortium, 2012; Mortazavi et al., 2013; Cheng et al., 2014; Yue et al, 2014). Also, SOMs were used to analyze co-binding of 208 TFs in HepG2 cells (Patridge et al., 2020). The Linked SOM method combines the clustering of multiple SOMs, each SOM built with a different type of data, into one analysis. By adapting the Linked SOM method to the data available for *Xenopus* mesendoderm development, we have developed a new approach that can be used to understand the GRN underlying germ layer specification.

To apply the Linked SOM method, two RNA SOMs were created by training on 95 transcriptomic RNA-seq data sets to identify 2 distinct sets of metaclusters that capture gene expression profiles co-varying across different experimental conditions, including spatial dissections. Similarly, a DNA SOM was generated by first creating >731,000 genome partitions based on co-varying TF binding, histone marking, and ATAC-seq peaks. These partitions were, then, clustered over 63 ChIP/ATAC-seq experiments. Next, we linked these SOMs by associating the individual partitioned genomic regions within the DNA SOM metaclusters to transcription units contained within the RNA SOM metaclusters. Motif analysis of the spatial linked metaclusters spotted several TFs with potential targets that had significant spatial differences, which we were able to add to our network. The linked SOM was then tested to determine whether it can predict functionally relevant TF-DNA interactions in the *Xenopus* mesendoderm GRN by focusing on the Foxh1 TF. All 19 previously identified direct connections were successfully recovered (Charney et al., 2017a). More importantly, the linked SOM has identified previously unsuspected, but functional, TF-DNA interactions as well as predicting combinatorial TF-TF interactions in *Xenopus* mesendoderm formation. These inferred interactions were validated using reporter gene assays, supporting that the linked SOM approach is a valuable method to build mechanistic GRNs.

## Results

### Reconciling the known biology with evidence from high-throughput data

The scope of the problem of building GRNs with high-throughput genomic data can be illustrated by examining the known regulatory loci surrounding the gene encoding the Spemann organizer TF Gsc (Cho et al., 1991), which contains two regulatory elements near the promoter called the proximal and distal elements (Watabe et al., 1995), and a farther upstream element (Mochizuki et al., 2000). These DNA regions are bound and controlled by a small set of known mesendodermal TFs (Koide et al., 2005). However, recent ChIP-seq datasets highlight the possibility of the function of various maternal TFs (Nakamura et al., 2016; Charney et al., 2017b; Paraiso et al., 2019), organizer TFs (Yasuoka et al., 2014) and mesodermal TFs (Gentsch et al., 2013) in regulating *gsc* through these cis-regulatory modules (CRMs; Figure 1A). Further, these known elements represent the minority of peaks identified upstream of the *gsc* transcription start site, as at least four other as yet unstudied genomic regions are occupied by various combinations of TFs. The chromatin context within this region highlights the need for integrative analysis of genomic datasets.

**Figure 1.**
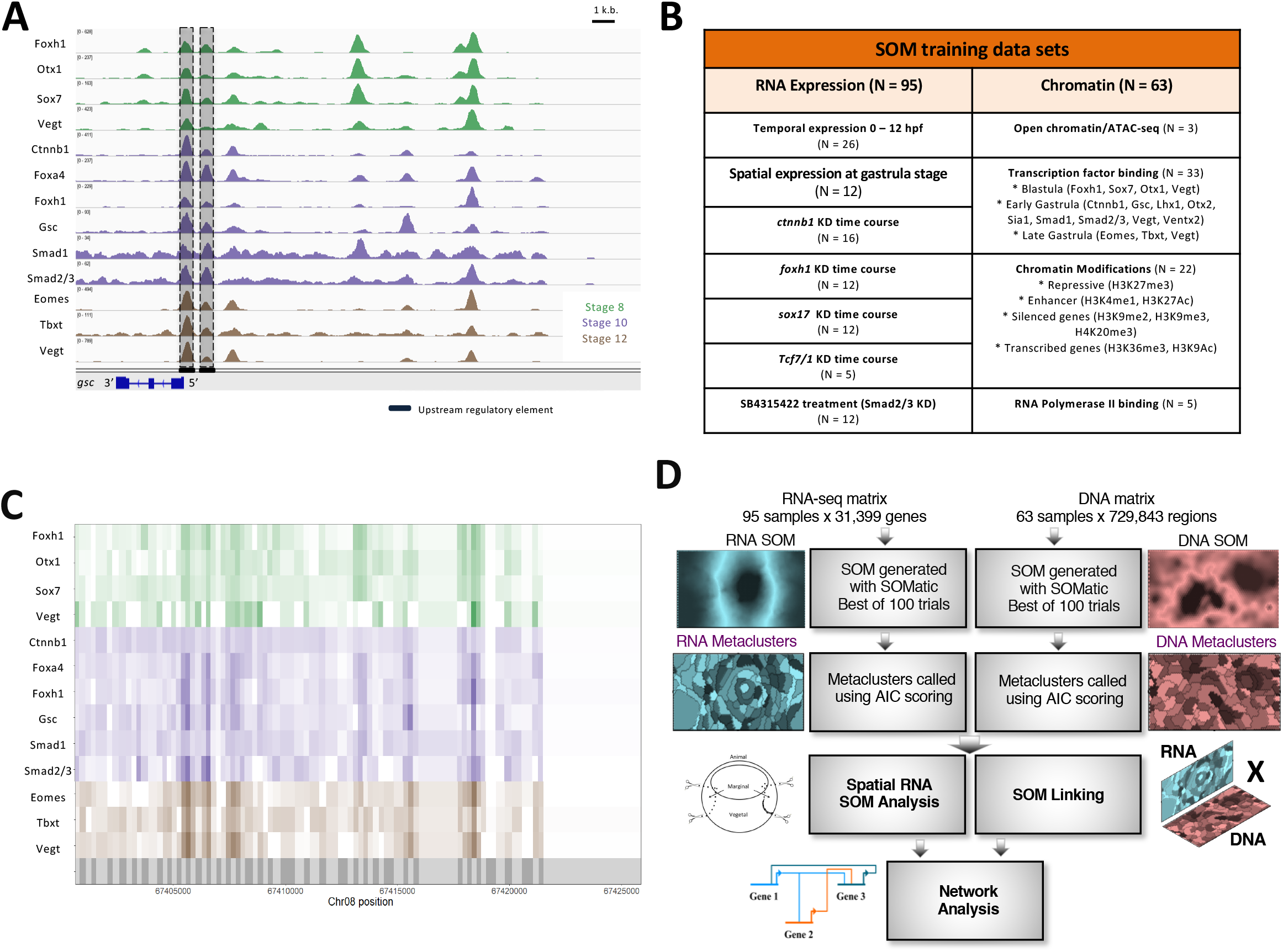
Using self-organizing maps to discover the Xenopus tropicalis mesendoderm regulatory network. **(A)** Genome browser view of transcription factor (TF) binding during early *Xenopus tropicalis* development. Shown are maternally-expressed (Foxh1, Otx1, Sox7, Vegt Ctnnb1, Smad1, and Smad2/3) and zygotically-expressed (Foxa4, Gsc, Eomes, Tbxt and Vegt) TF binding in the gsc gene locus. Shaded are the well characterized proximal, distal, and upstream cis-regulatory elements, which are associated with TF binding. Further upstream are binding sites in possibly unexplored cis-regulatory regions **(B)** Datasets used in this analysis. There are many different experiments targeting several wild-type and MO-injected embryos at developmental stages important for mesendodermal development. **(C)** The *X. tropicalis* genome is partitioned using ChIP-seq and ATAC-seq peak locations. Each partition is then assigned ChIP-seq and ATAC-seq signal quantified as reads per kilobase per million (RPKM) for all chromatin datasets. **(D)** The RNA-seq and ChIP-seq/ATAC-seq data sets were each converted into training matrices and were clustered using SOM metaclustering using SOMatic. These clusters were then linked using the SOM Linking tool within SOMatic. The pair-wise linked metaclusters (LM), and spatial SOM data were mined for regulatory connections and built into networks.

### Strategy for integration of different highly-dimensional genomic data types

To investigate *Xenopus tropicalis* mesendodermal gene regulation, we assembled a highly dimensional data set of 95 RNA-seq and 63 ChIP-seq/ATAC-seq experiments (Figure 1B; see Key Resources Table). The chromatin datasets include the openness of the loci (ATAC-seq), transcription factor binding, epigenetic modifications (e.g. enhancer and polycomb histone marks), and RNA Polymerase II association. The transcriptomic datasets include spatial expression (e.g. ectoderm, mesoderm and endoderm) and temporal expression at relevant developmental stages. These datasets also include knockdown of critical TFs (e.g. Sox17 and Foxh1) and signaling pathways (e.g. Nodal and Wnt signaling), which regulate mesendoderm formation. These data were individually analyzed and collected into two large matrices for unsupervised learning (see STAR Methods). For the RNA-seq experiments, gene expression was quantified in transcripts per million (TPM) for each experiment. For the ChIP/ATAC-based experiments, in order to annotate distinct DNA regions across the whole *Xenopus* genome in a similar fashion to ChromHMM (Ernst and Kellis, 2012), we first partitioned the genome using peak calls identified from these chromatin datasets. Then, we calculated reads per kilobase per million (RPKM) signal for each experiment within each of these partitions. As an example, compare the conversion of the ChIP-seq peaks (Figure 1A) to the RPKM ChIP-seq signal (Figure 1C) in the *gsc* locus. These normalized tracks properly transform the raw signal into a form that our downstream unsupervised neural networks can accept to perform clustering. We also note the region furthest upstream of *gsc*, which does not contain any peaks (potentially quiescent regions).

Next, we implemented a strategy for highly-dimensional data integration through linked SOMs (Jansen et al., 2019) (Figure 1D). This involved performing unsupervised learning through SOMs on each type of data (i.e. RNA-seq data and chromatin data) separately, followed by metaclustering to generate a separate RNA SOM and DNA SOM. For the RNA SOM, genes are clustered in terms of similarities in expression: spatially, temporally, and by effects of perturbation. For the DNA SOM, potential cis regulatory information, the DNA partitions are clustered based on similarities in DNA accessibility, histone modifications, and combinations of TFs bound. These RNA SOM and DNA SOM clusters are then compared to each other and linked such that clustered transcription units are associated with nearby clustered DNA partitions (i.e. clusters of DNA partitions are linked to the RNA cluster membership of their nearby genes). Additionally, separately created spatial RNA SOM clusters were incorporated into the network analysis (discussed further below). This combined approach allows for groups of cis-regulatory genomic regions to be linked to their nearby target gene expression profiles for further network analysis (Figure 1D).

### RNA SOM identifies gene expression modules

To identify different gene expression cohorts (modules) present during early *Xenopus* development, we performed unsupervised learning on the RNA-seq experimental data by training a SOM followed by metaclustering (see STAR Methods). Due to the experimental matrix being “dominated” by time course expression data, the trained map displayed a time course-dependent structure such that genes that have similar temporal profiles such as *gsc*, *nodal1, lhx1, osr1, hhex*, and *osr2* being located in SOM units in one general area of the 2D map, whereas genes that tend to peak earlier, such as *nodal, nodal2*, and *sia1*, were in SOM units in another section (Figure S1A) (Owens et al., 2016). On the other hand, while comparisons between gene expression knockdowns using antisense morpholino oligonucleotides (MOs) and their controls were a minority in this dataset (Figure 1B), they do show local differences on the 2D maps across adjacent metaclusters (Figure S1B). Thus, the metaclustering of the units of the map had the capacity to capture all of these differences and the data are available at https://hpc.oit.uci.edu/~csjansen/SOMs/xenoRNA.60x90.v18/.

In all, we recovered 84 distinct RNA SOM metaclusters that capture the different gene expression profiles present in the included data (Figure 2A, Figure S1B, Table S1). Genes that share similar expression profiles across all experiments such as dorsal mesendodermal genes activated during midblastula stage, including *nodal, nodal2*, and *sia2*, clustered together in metacluster R82. Organizer genes *gsc* and *hhex*, which showed transient zygotic expression peaking at stage 10 clustered together in metacluster R11 (Figure 2B). Meanwhile, genes in metacluster R76 (Figure 2C), which include *foxa2* and *gli3*, did not become highly expressed until mid-gastrula stage 11 and steadily increased until the end of the experimental measurements (stage 13). In addition to spatio-temporal expression, genes within each metacluster showed distinct responses to perturbation experiments (Figure 2C). For example, the 110 genes in metacluster R11 had similar responses to multiple MO perturbation and temporal conditions. Notably, the genes in this group were down-regulated in stage 10 after inhibiting Foxh1 expression (Foxh1 MO experiment), whereas the 728 genes in R23 and the 527 in R72, were up-regulated. To further show that each metacluster is distinct, we performed Gene Ontology enrichment analysis on each (Figure 2D). Metacluster R11 contained genes with functions related to dorsal/ventral patterning and cell fate, whereas metacluster R72 had genes related to cell proliferation and R23 had genes associated with post-translational modifications. See Table S2 for other GO term analysis of metaclusters. The differences in biological functions and properties among these GO term lists suggests that the RNA SOM distinguishes sets of genes based on their expression behaviors under different conditions.

**Figure 2.**
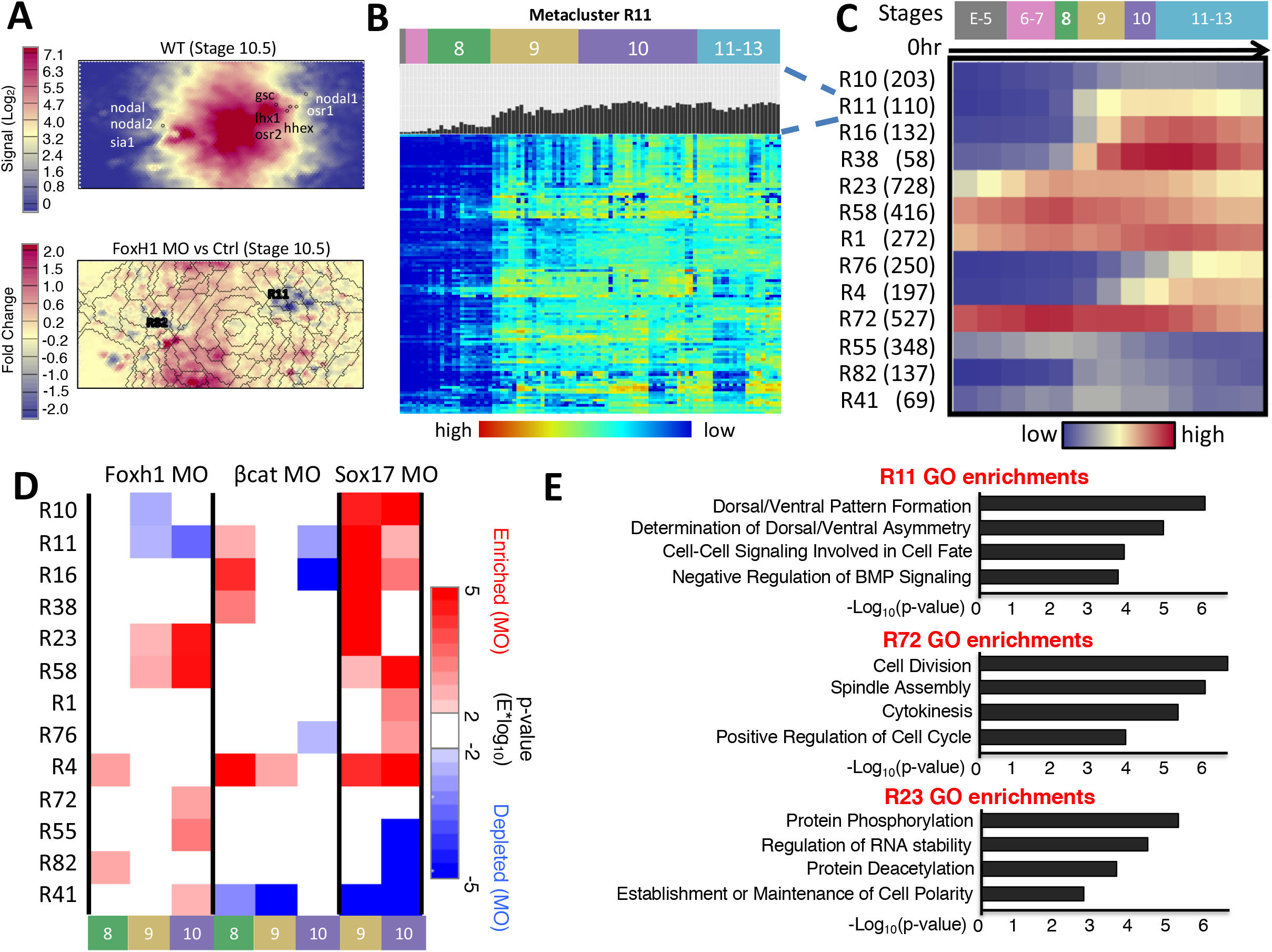
RNA-seq SOM metaclustering reveals developmental gene modules that contain similarly regulated genes. **(A)** Metaclusters containing genes from the core mesendoderm network show unique temporal dynamics during development. *nodal, nodal2* and *sia* are grouped left and *gsc, nodal1, lhx, osr2* are grouped right (top). 2D metaclustering with overlaid metacluster boundaries shows the genes that are up and down-regulated upon Foxh1 MO knockdown (bottom). **B)** Each metacluster is filled with genes with a similar expression profile, for example a heatmap of the genes in metacluster 11 is shown. (**C**). Heatmap of average temporal expression profiles of genes belonging to 13 RNA metaclusters. Parentheses after RNA metaclusters indicate number of genes in each RNA metacluster. **(D)** Two-tailed hypothesis analysis applied on gene metaclusters after subtracting out control experiments. Each metacluster responded to each MO experiment differently at different time points. **(E)** GO term enrichments for genes within three example RNA SOM metaclusters. Each metacluster had unique functional enrichments supporting the coherence of these clusters.

### DNA SOM identifies chromatin states and combinatorial TF binding

To prepare the data from the collected ChIP-seq/ATAC-seq experiments for machine learning, we separated the *Xenopus tropicalis* genome into 731,726 genome partitions using called peaks from each experiment (Figure 1C, Figure S3) and computed RPKMs for each experiment over these regions. We then performed unsupervised learning on this matrix with a SOM, and further metaclustering identified 88 distinct DNA profiles present in the data (Figure 3A, Figure S3). Like ChromHMM, these clustered partitions are differentiated by histone marks according to different chromatin states such as H3K4me1-marked active or primed enhancers (metacluster D71 and D58; Figure 3A) (Heintzman, et al., 2007; Creyghton et al., 2010; Buenrostro et al., 2013), H3K9me2/3- and H4K20me3-marked heterochromatic regions (metacluster D72 and D29; Figure S3)(Schotta et al., 2004) and unmodified regions (metacluster D9; Figure S3) (Hontelez et al., 2015). Additionally, the hierarchical clustering over these metacluster profiles Polycomb repressive H3K27me3 marked regions (Cao et al., 2002) are separated from other chromatin marks in metacluster D45 and D84 (Figure S3). Similarly, promoter regions marked by H3K4me3 ChIP-seq signals clustered together (D28 and D87; Figure S3) (Santos-Rosa et al., 2002). Interestingly, metacluster D51 has a strong H3K27me3 and H3K4me1 signal, which indicated that this metacluster contains inactive promoters and putative poised enhancers, whereas metacluster D77 replaced the H3K27me3 signal with H3K27ac, which indicated active promoters and Ep300-positive enhancers.

**Figure 3.**
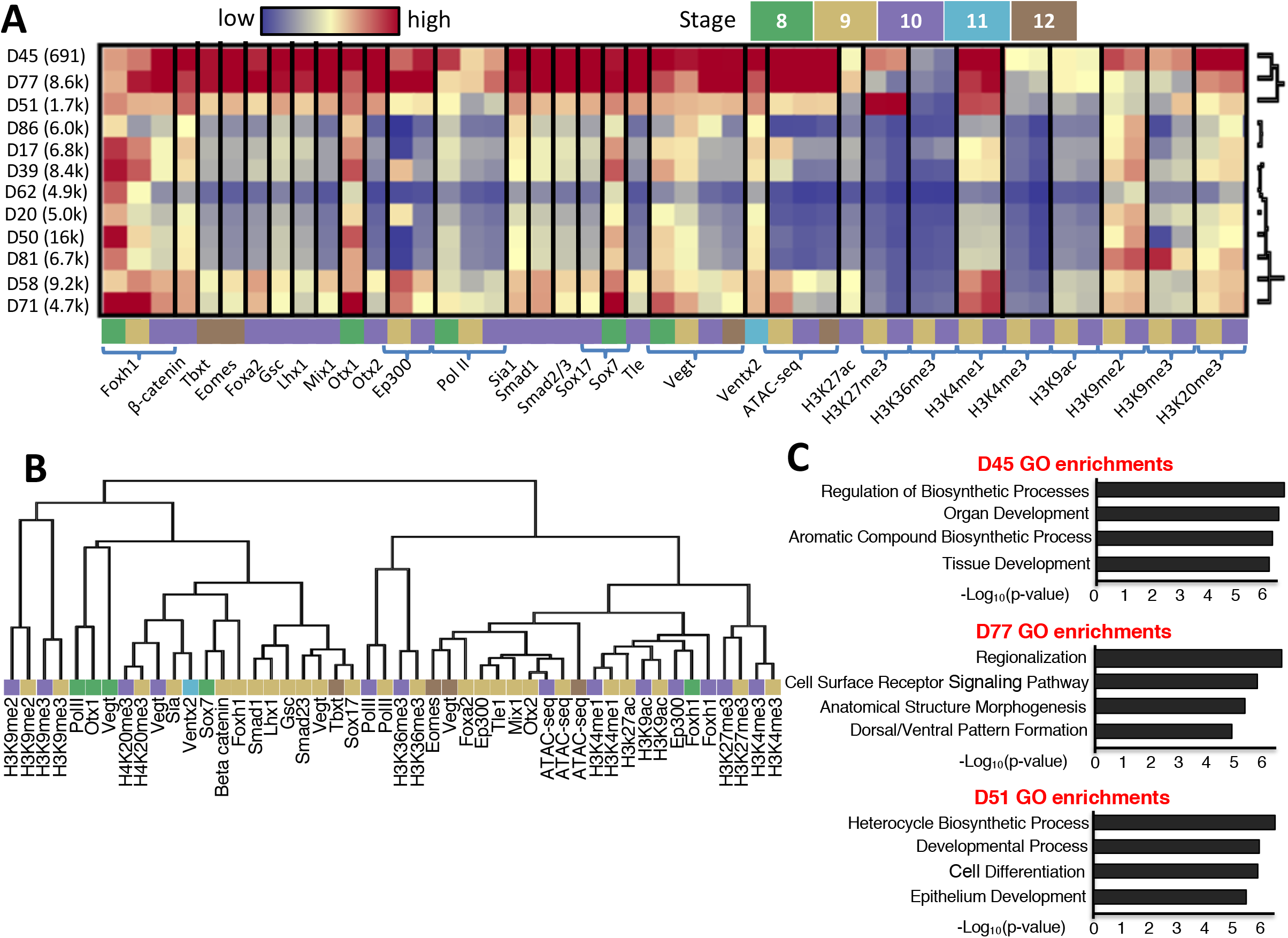
Self-organizing map-based clustering shows Foxh1 co-binding and functional gene modules during gastrulation. **(A)** Heatmap of Foxh1 ChIP-enriched metaclusters that visualizes the different patterns of co-regulation present in Foxh1-bound CRMs. **(B)** Experiment hierarchy of ATAC/ChIP-seq data after metacluster correction. The developmental stages of each experiment are indicated by the same color coding as (a). **(C)** GO term enrichments for genes nearby genome regions within three example ATAC/ChIP SOM metaclusters.

Next, we searched within enhancer-marked regions and visualized interactions of known TF co-bindings via 2D-SOM. For example, previously, we have shown that the maternally-expressed endodermal TFs Otx1, Vegt, and Foxh1 can co-bind CRMs, and Otx1 and Vegt synergistically activate endodermal gene expression during cleavage to early blastula stage 8 (Paraiso et al., 2019). In this analysis, Figure S1C highlights that Otx1, Vegt, and Foxh1 ChIP-seq data showed considerable overlap and simultaneous enrichment in metaclusters D77, D71, and D50 (Figure 3A). Secondly, there are significant metacluster overlaps (D20, D39, D58, D71, D77, D45, and D51) between Ep300 and Foxh1 at stage 9 (Figure S1D), indicating a close association between these two factors. Thirdly, unlike the early binding of Foxh1 during blastula stages 8-9, Foxh1 binding during early gastrula stage 10 is enriched near dorsal mesendodermal genes and is associated with the Nodal co-factor Smad2/3 binding as seen in D77 (Chiu et al., 2014; Charney et al., 2017b). Consistent with this observation, Foxh1 binding and Smad2/3 binding were highly correlated as shown by an extensive overlapping heatmap, whereas a heatmap representing Foxh1 binding during blastula stage only partially overlapped with the Smad2/3 heatmap (Figure S1D, Figure S3). The fact that metaclusters, D39, D58, and D71 are free of Smad2/3 but associate with Ep300 indicates that many of Foxh1 bound regions have a Nodal signaling independent activity. Lastly, at late gastrula stage 12, T-box TFs Eomes, Vegt and Tbxt (Brachyury), which co-regulate mesodermal gene expression (Gentsch et al., 2013), showed similar binding profiles among metaclusters D38 and D54 (Figure S1E, S3). The clustering in Figure S3 shows a close interaction between Vegt at stage 10.5 and Tbxt at stage 12. Tbxt at stage 12 is expressed in mesoderm originating from the organizer tissue and the circumblastoporal collar, whereas Vegt at stage 10.5 is detected in both mesoderm and endoderm. Thus, the close interaction between Tbxt and Vegt appears to be from a mesoderm origin. Vegt and Eomes may have clustered together due to sharing a highly conserved T-box amino acid sequence (Conlon et al., 2001)(Figure S3). However, the binding profile of Tbxt, which has a slightly divergent T-box amino acid sequence did not cluster as well with its closely-related family member TFs (Sebé-Pedrós el al., 2013) at stage 12. Perhaps, this represents a situation in which Vegt and Eomes regulate endodermal genes, while Tbxt regulates mesodermal genes. These analyses illustrate the advantage of presenting the ChIP-seq data with a SOM analysis to visually inspect TF-TF interactions and uncovering functional differences of closely-related TFs.

Outside known interactions, we find some surprising combinations of TF binding. For example, there is a substantial overlap between the Foxh1 and Gsc SOM maps from the stage 10.5 gastrula (Figure S1D). Interaction between Gsc and Foxh1 has not been well documented in the past, but there is evidence that they directly interact and regulate the expression of the *mix1* gene, encoding a TF involved in endoderm development (Izzi et al., 2007). Our SOM results suggest that such an interaction may be more wide-spread during *Xenopus* mesendoderm specification. Additionally, the binding of mesodermal regulator Tbxt (Smith et al., 1991) at stage 12 and the endodermal regulator Sox17 (Hudson et al., 1997; Mukherjee et al., 2020) at stage 10.5 correlated with each other, more so than with the binding of other TFs (Figure S1E, Figure S3). This finding is interesting as Sox17 is generally considered to be an endodermally expressed gene, whereas Tbxt is mesodermally expressed. If Sox17 remains bound to these regions until stage 12, this could indicate either a competitive or an independent interaction between Tbxt and Sox17 at that stage to generate distinct mesodermal and endodermal lineages. In support of the latter, the expression patterns of Tbxt and Sox17 are also mutually exclusive in mice (Lolas et al., 2014), indicating that may represent a conserved mutual exclusion mechanism regulating mesoderm and endoderm development. Lastly, at early gastrula stage 10, the binding of dorsal mesendoderm factors Ctnnb1 (β-catenin; Wnt signaling transducer) (Nakamura et al., 2016; Heasman et al., 1994) and Foxh1 (Nodal signaling cofactor) (Chen et al., 1996) clusters with the ventral specifying TF Smad1 (Bmp signaling transducer) (Graff et al., 1996; Afouda et al., 2020) (Figure S1F, Figure S3). Some of these CRMs that show interactions with Wnt, Bmp, and Nodal signaling pathways may represent the nodes critical in controlling the formation of the dorsal-ventral axis during early embryogenesis. These newly identified combinatorial interactions of TFs underline the usefulness of SOM analysis and would be the topic of further research.

### Distinct genes and consensus DNA binding motif profiles are associated with different DNA SOM metaclusters

To further characterize DNA metaclusters, we performed Gene Ontology enrichment analysis on the nearest genes of these regions (Figure 3B) (Table S4 for the full list). Embryonic processes correlated with the gene set associated with DNA regions in metacluster D45 are linked to organ and tissue development, while those near metacluster D77 are associated more specifically with morphogenesis and patterning. Additionally, the genes near regions in metacluster D51 are enriched for GO terms associated with cellular and developmental processes. D51’s profile of strong H3K27me3 and H3K4me1 marks with weak ATAC-seq and Ep300 signals. When matched with the RNA metaclusters, these genes were highly enriched in R4, R16, and R76 (See Figure 2A for these profiles) which were all characterized by expression at later timepoints. The GO analysis, thus, indicated the genome segments in these DNA metaclusters are used in different transcriptional programs, and so, require differential gene regulation.

Further, in order to identify the TFs that may control the expression of genes with these distinct metaclusters (D45, D51, and D77), we performed consensus DNA binding motif analysis on each metacluster using the HOCOMOCO v11 human motif database (Kulakovskiy et al., 2017) with a q-value of 0.1. After removing the shared motifs among the metaclusters, 63 unique TF motifs to metacluster D45 were recovered, such as Smad2/3, Sox7, and Ventx. These are well known TFs involved in mesendoderm development (Lagna et al., 1996; Ault et al., 1996; Schmidt et al., 1996; Zhang et al., 2005). Metacluster D77 contained 56 unique TF motifs, including Foxa2 and Tcf3 (also known as E2a), which are important in the regionalization of mesendoderm (Zorn and Wells, 2009; Wills and Baker, 2015). Finally, 37 TF motifs including Gata6, which is important for endoderm development (Afouda et al., 2005), are found in metacluster D51. D51 also includes the Tead1 motif, which is interesting because it is a known repressor in stem cells (Maeda et al., 2002) and the regions in D51 are also decorated with the repressive H3K27me3 mark. Based on these analyses, we concluded that the DNA SOM clustering managed to separate the genome partitions into groups of genes with potentially different biological functions.

### A spatial RNA SOM discovers independent spatio-temporal gene modules

Next, we wished to explore the spatial RNA-seq data in more detail. Unlike the morpholino (MO) knockdown data, which was successfully incorporated into the SOM metaclustering (Figure 2D), genes in the full RNA SOM did not separate on the map based on their spatial expression profiles. This was due to the full RNA SOM being too focused on the temporal data provided, and so, we decided to perform a parallel analysis to provide further insights. For this, we trained a separate SOM, and performed metaclustering, based on just the spatial RNA data from dissected early gastrula (stage 10.5) tissues (Blitz et al., 2017). This analysis provided an excellent separation of genes based on their spatial expression (Figure 4A), and the metaclustering separation followed the differential areas of the map well. Spatial metaclusters sR9, sR8, and sR1 especially had quite visible differential spatial gene expression, and sR15 and sR8 showed differential gene expression when the average fold change from each experiment to the whole embryo for each metacluster was plotted (Figure 4B). To show statistical significance of this differential expression, the hypothesis tool in SOMatic was used to find that sR1, 6, 8, 9, 12, and 15 were statistically significant in difference from whole embryo expression levels (Figure 4C). sR8, 15 and 9 were enriched in the endoderm (vegetal pole), and sR12, 1 and 6 were enriched in both mesoderm (dorsal, ventral, and lateral marginal zones) and ectoderm (animal cap).

**Figure 4.**
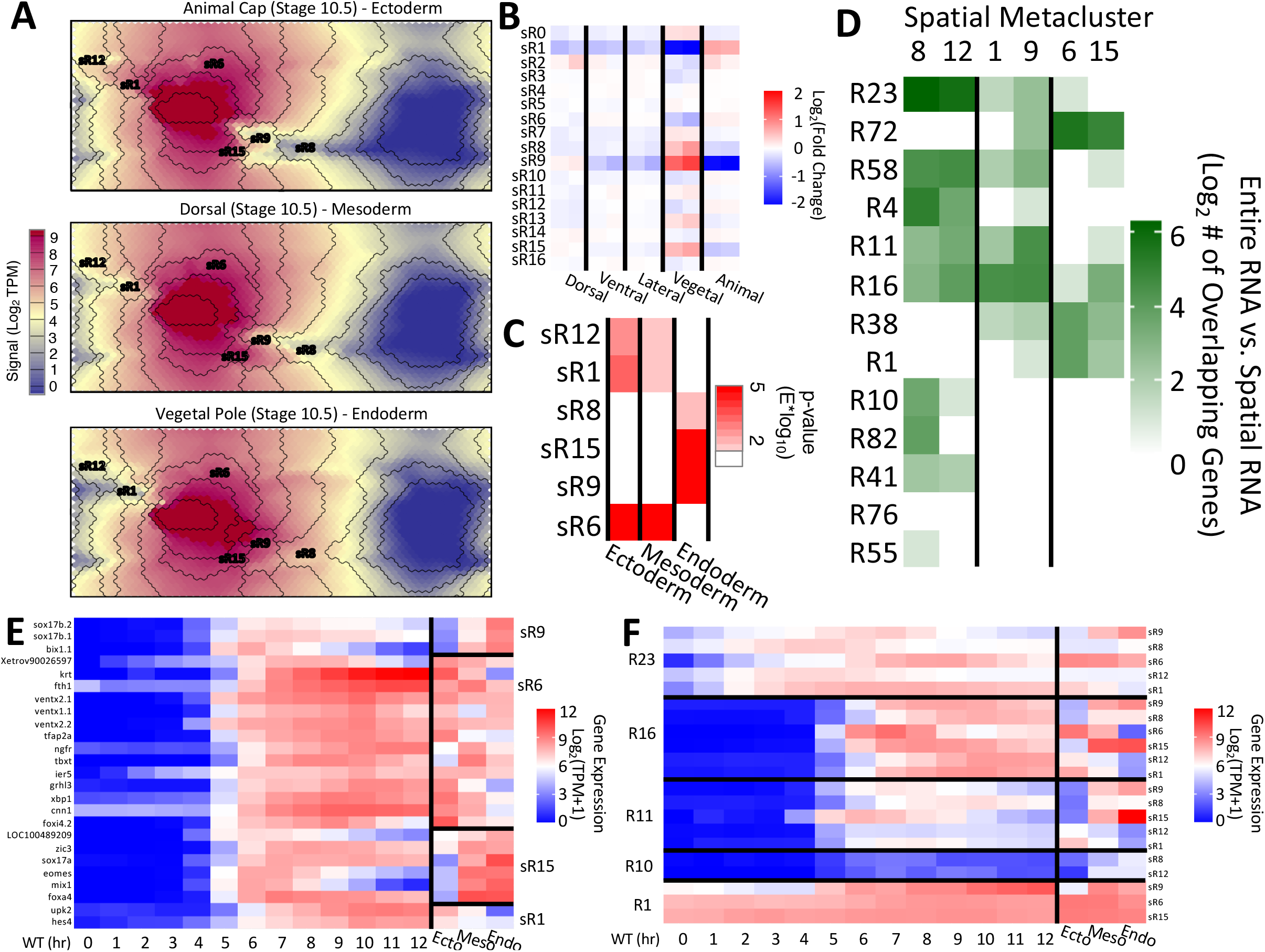
RNA metaclusters can be further segregated by spatial RNA SOM. **(A)** SOM slices from the spatial RNA SOM analysis corresponding to RNAs from the animal, dorsal, and vegetal tissue fragments with overlaid spatial metacluster boundaries. Some important spatial metacluster locations are noted. **(B)** Heatmap of the fold change of spatial metaclusters compared to the whole embryo experiments. **(C)** Heatmap of statistical difference between gene expression in each tissue and the whole embryo. 6 spatial metaclusters (sR) showed statistically significant differences in ectoderm/mesoderm or in endoderm. **(D)** Joint membership of genes in spatial RNA metaclusters and RNA metaclusters from the full RNA data set. Rows and columns are hierarchically clustered. **(E)** Temporal (from wild-type) and spatial gene expression profiles for genes in both sR 9, 6, 15, and 1 and R38. **(F)** Average temporal and spatial gene expression profiles for genes in both R23, R16, R11, R10, or R1, based on spatial metaclusters.

To further explore these sets of genes, we overlapped them (Figure 4D) with the full RNA SOM clustering shown in Figure 2C. Hierarchically clustering the spatial RNA metaclusters based on gene overlap showed 3 separate spatial metacluster groupings, sR8/sR12, sR1/sR9, sR6/15, that had similar overlaps with the full RNA SOM. Interestingly, each group contained 1 metacluster significantly enriched in the endoderm and the other enriched in the ectodermal and mesodermal experiments (e.g. see Figure 4C, compare sR8 and sR12), and each grouping had a specific set of full RNA metaclusters (temporal profiles) that it overlapped with. These observations suggested that there might be sets of potential spatial-specific TFs activated simultaneously in different parts of the embryo that bring about the spatial gene patterns we see in the developing embryo.

To ensure that this observation was not an artifact of the clustering method, we plotted the raw profiles of multiple genes in 1 full RNA metacluster (R38) and classified those genes by their eventual membership in the differential spatial RNA metaclusters (Figure 4E). Based on the time course data, these genes are activated at about 5 hours of development, and they each have very different spatial profiles (panel E, right, ecto, meso, endo). We also plotted the average profiles of each of the genes in each of the metacluster overlaps (Figure 4F). As expected, the temporal profiles of the genes match in each of the full RNA SOM metaclusters. However, we noted significant spatial differences. This prompted us to explore the regulatory elements near these genes to identify the spatial-specific TFs that are driving this behavior

### Multi-omic data integration of ChIP/ATAC SOM and Spatial RNA SOM provides direction of transcription output

Previously, we developed the linked SOM method specifically to integrate scRNA-seq and scATAC-seq datasets (Jansen et al., 2019). Metaclusters of a scRNA-seq SOM were linked to a scATAC-seq SOM to build sets of genome regions that had similar scATAC-seq profiles near genes with similar scRNA-seq profiles. We determined that this linked SOM approach could be implemented similarly with the spatial *Xenopus* RNA-seq and the abundance of ChIP/ATAC-seq data to uncover the TFs that drive the observed spatial patterns.

Our goal here is to identify specific TF motifs that are enriched among genes that are expressed in specific regions of embryos. We applied the linked SOM approach to the spatial RNA SOM and DNA SOM and generated a linkage between the 16 spatial RNA and the 88 DNA metaclusters, resulting in 1408 (16 x 88) linked metaclusters (LMs). A motif search using Find Individual Motif Occurrences (FIMO) was performed on each LM separately using the HOCOMOCOv12 human motif database and motifs that were specifically enriched in a subset of LMs were identified. Of these motifs, we focused on those that appeared near genes in the six differential spatial metaclusters (Figure 4C) forming 3 groups: sR8/sR12, sR15/sR6, sR9/sR1 (Figure 4D). For each pairing, any motifs that appeared in the union of the spatial metaclusters were filtered out, and motifs that were specific to one metacluster were retained (Supp Fig 4A). To further enrich for TFs with targets showing spatial expression, we searched for the motifs that were shared in at least 2 of the sets (Supp Fig 4B). Then, we plotted the temporal/spatial expression of 20 candidate TFs that could bind to these motifs (Figure 5A&B).

**Figure 5.**
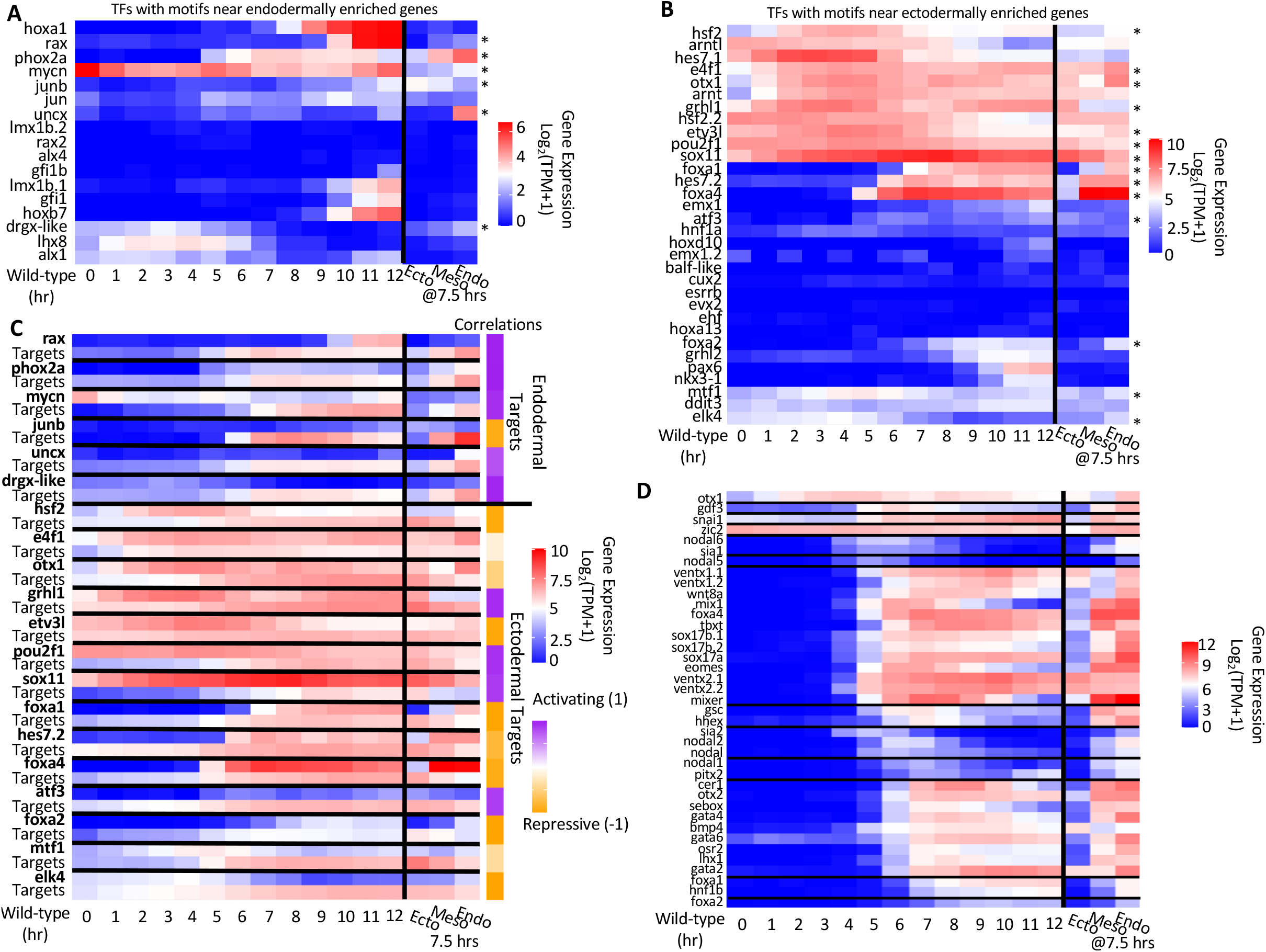
Spatial metaclustering assists in identifying candidate TFs for Xenopus mesendodermal differentiation. **(A-B)** Temporal and spatial gene expression profiles of TFs with motifs found near endodermally (A) or ectodermally (B) enriched genes. Asterisks indicate TFs that show distinct spatial expression patterns. **(C)** Temporal and spatial gene expression profiles for spatially differential TFs (bold) matched with the average gene expression profile of their predicted targets (Targets). Correlations were calculated by comparing their spatial gene expression profiles. **(D)** The temporal and spatial gene expression profiles of genes known to be important in *Xenopus* mesendodermal development, separated by RNA metacluster.

Among those, we selected the TFs that showed a significant (qvalue < .05) differential spatial expression. From the motif set near endodermally-enriched genes, 6 TFs showed significant differential spatial expression (Figure 5A, asterisks), whereas, from the motif set near ectodermally-enriched genes, we found 14 TFs (Figure 5B, asterisks). Figure 5C shows the temporal/spatial expression profiles of these TFs alongside the average expression of the predicted targets of these TFs in the differential spatial metaclusters. By computing the correlation of the spatial signal from the TFs and the expression levels of their predicted targets, we predicted the direction of transcriptional output - potential activating or repressing roles of these TFs. For example, in ectoderm where *foxa1* and *foxa4* expression is low relative to endoderm, target gene expression levels are high in ectoderm. This suggests that Foxa1 and Foxa4 had a strong negative correlation between their spatial expression and their potential targets, indicating that they have a major role in repressing mesodermal and ectodermal fates.

One interesting observation from this plot is that the majority of predicted spatial regulators of endodermal targets are activators, whereas, for the ectodermal targets, most regulators are repressive in nature. This is consistent with the view that mesendodermal cells are induced from pluripotent cells that differentiate via an ectodermal default path. For cells to differentiate from a ectodermal to a mesendodermal state, certain ectodermally (animally)-expressed genes need to be reduced in expression (through endodermally/vegetally-expressed repressors) and other genes need to be expressed (through endodermally-expressed activators). Of the 14 ectodermal TFs, only 4, Ghrl1, Pou2f1, Sox11, and Atf3, had a positive spatial correlation with their targets. Some of these genes are known activators in ectodermal tissues in other organisms (Edgar et al., 2013). Sox11 is a positive regulator of neuronal differentiation in frogs, chick, and mouse (Bergsland et al., 2006, Lin et al., 2010, Chen et al., 2016). Pou2f1 (previously known as Oct1) is an activator that is expressed in a wide variety of cell types, including in ectodermal cell lineages in *Xenopus laevis* (Veenstra et al., 1995).

Of more interest to this work, there were 10 TFs with motifs near ectodermally-expressed genes that were marked as repressive as their expression was significantly higher in the endoderm. Some of these were already included in the core mesendodermal network, such as *foxa1, foxa2, foxa4*, and *otx1*, with others being new potential additions. The data support the notion that these TFs have a repressive role to suppress unwanted ectodermal gene expression in the endoderm. Of the 6 new TFs, only 2 are expressed at high enough levels at stage 10 to be considered for being added to the core network: *hsf2* and *hes7.2*. Additionally, there are 3 TFs with motifs that were found near endodermal genes with a high enough gene expression at stage 10 to be considered as well: These are *phox2a, mycn*, and *uncx*. Each of these genes have similar temporal/spatial profiles to the genes from the core mesendodermal network (Figure 5D) and were included in the downstream network analysis.

### Generation of a comprehensive mesendodermal GRN using multi-omic data integration

In our hand-curated mesendoderm GRN (Charney et al., 2017a), a bipartite criterion was used to determine direct TF regulation, whereby a gene was considered a likely TF target if its expression is both affected by the perturbation of the TF and if the CRMs near the gene show physical association with the TF. This work required a large investment in manpower and effort, and yet the network was incomplete. With the success of the linked SOM method on finding specific motifs from the spatial RNA and DNA data, we moved to implementing the approach on the full RNA SOM and DNA SOM. This generated a linkage between the 84 RNA metaclusters and the 88 DNA metaclusters, resulting in 7,392 (84 x 88) linked metaclusters (LMs).

Unlike the spatial/DNA linked SOM analysis, we were interested in using a set of more specific motifs for known maternal and signaling factors, and as such, we utilized ChIP experiments to build a *Xenopus*-specific DNA binding motif database of Eomes, Foxa2, Foxh1, Gsc, Mix1, Otx1, Otx2, Smad2/3, Sox17, Sox7, and Vegt (Table S5) and motifs for human Tcf7l1/2. When we scanned each LM for these motifs using FIMO with a q-value of 0.1, we received a set of 271,736 total significant motif instances. Of these, the largest portion belonged to Foxh1 with 134,238 detected motif instances. These initial motif lists were again filtered by LM motif density (see STAR Methods) to find significantly (pvalue < 0.05) represented motifs in each LM, which reduced the overall number to 201,157 with 118,722 belonging to Foxh1.

Next, we developed a filtering strategy to focus on the targets active at the developmental time of interest, starting with Foxh1 targets. Limiting the Foxh1 motif instances to those in DNA metaclusters with Foxh1 ChIP-signal in the 75th percentile near genes in RNA metaclusters with significant gene expression (> 1 TPM) in stages 8-10.5 reduced the number further to 117,253. This small reduction shows that the motif analysis was mostly concordant with the ChIP signal, even before filtering, suggesting that most of the 118,722 genome regions with an identified Foxh1 motif were actually bound by Foxh1. Next, to ensure that we only analyzed active Foxh1 binding sites, we incorporated ChIP-seq/ATAC-seq metaclusters that have an enriched Ep300 signal at stage 9. Application of this filter dramatically dropped the list of potential functional FoxH1 motifs from 118,722 to 26,445 and reduced the number of predicted target genes from 12,831 to 6,717.

To assess the quality of our GRN, we sought to estimate the false positive rate (FPR) for predicted Foxh1 targets. Since a set of true negative gene targets does not exist, we built a list of likely true negative targets for Foxh1 by calling significantly un-changing genes from each of the Foxh1 MO experiments (stages 8, 9, and 10) with DEseq2 (Love et al., 2014) and intersected the lists (Table S6). Of the 5,864 likely true negative target genes, 696 were found within our set of potential targets. This gave the analysis an 11.9% FPR (10.3% FDR), which we deemed acceptable.

To further focus the network, we employed additional constraints by selecting RNA metaclusters that contained genes that regulate gastrulation (Charney et al., 2017a), thereby filtering to 11,295 Foxh1 motifs located near 2,747 unique genes (Table S7 for full table). Next, we filtered out genes that did not encode TFs or growth factors from our previous works. After this process, 1,492 Foxh1 functional motifs were predicted to be near 242 TFs, and all genes from the curated core mesendodermal network (Charney et al., 2017a) remained in the list of 242 (Table S8). This final network includes 2,725 predicted connections for all 12 of our ChIPed TFs with 321 total targets. Finally, we visualized known and predicted network connections of the 36 targets of Foxh1, Sox17, Tcf7l1, Vegt, and Smad2/3 that were present in the core mesendodermal network (Charney et al., 2017a) (Figure 6A; Table 1). Of these, 17 connections for Foxh1, 11 for Sox17, 8 for Tcf7l1, 5 for Vegt, and 2 for Smad2/3 were new to this analysis, which does not include other new connections to the new members of the network. These new potential TF/gene connections inform us what other TFs impinge on mesendoderm GRN and thus should improve our understanding of the regulatory processes behind the determination of mesendodermal cell states.

**Figure 6.**
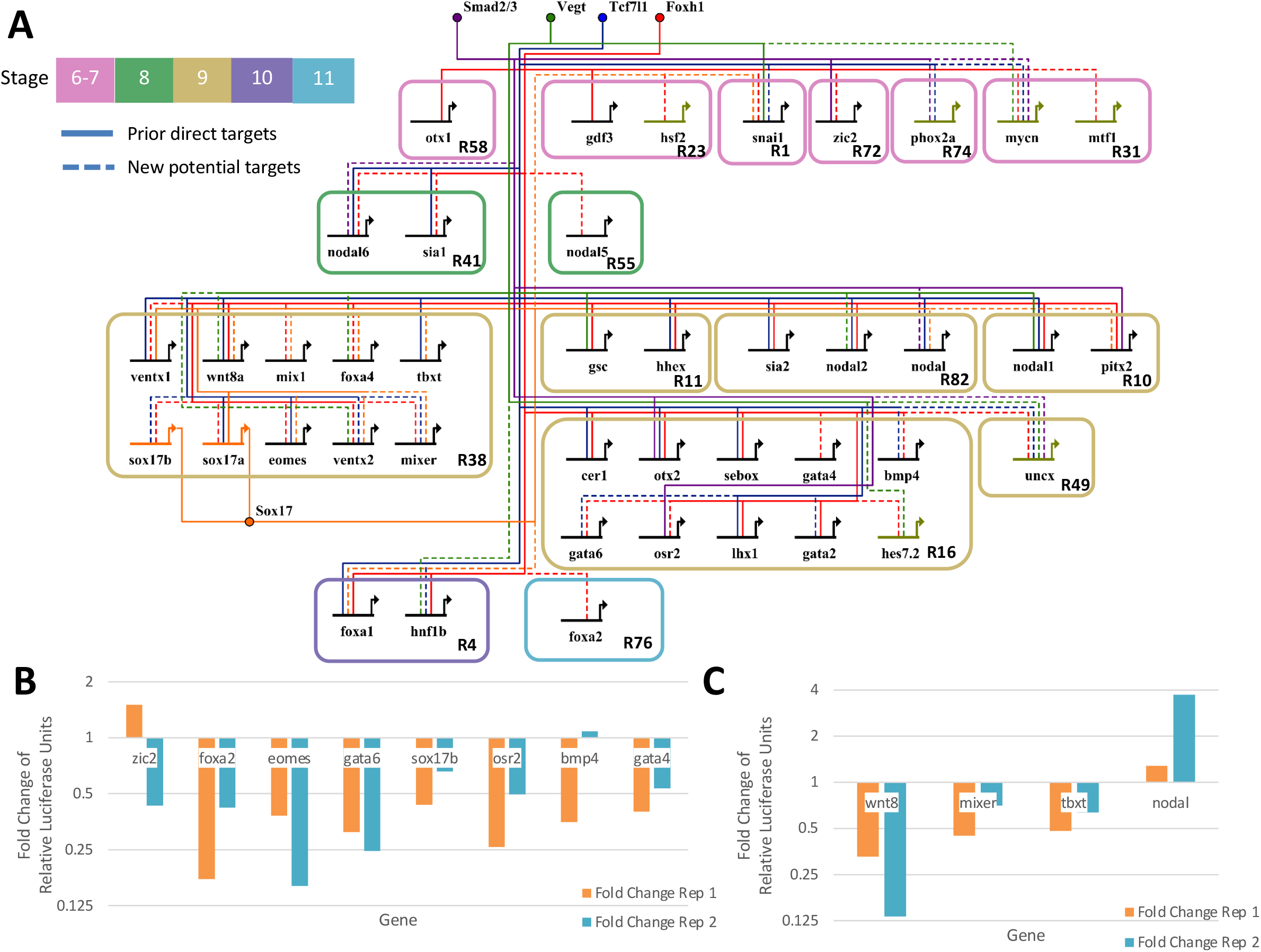
Gene regulatory network centered on the activity of Tcf7l1, Sox17, Vegt, Smad2/3, and Foxh1. **(A)** By linking the ChIP/ATAC-seq SOM metaclusters representing active cis-regulatory regions to RNA-seq SOM metaclusters, a gene regulatory network was generated. The active cis-regulatory regions were identified based on the enrichment of their respective TFs, enrichment of Ep300 signal, and DNA binding motif presence. Shown are literature identified targets (“prior direct targets”) which were validated by the Linked SOM pipeline and potential new connections (“new potential targets”) from the same pipeline. Note that only a subset of targets is shown and the network is focused only on TF and signaling molecule targets. **(B)** Fold change of relative luciferase units in Log scale after foxh1 MO injection. The luciferase activity was normalized to Renilla signal from pRL-SV40. Each of these shows enhancer activity, except *zic2* and *bmp4* being inconclusive. **(C)** Fold change of relative luciferase units after Sox17 MO injection in Log scale normalized to Renilla signal. Each of these shows enhancer activity.

**Table 1.**
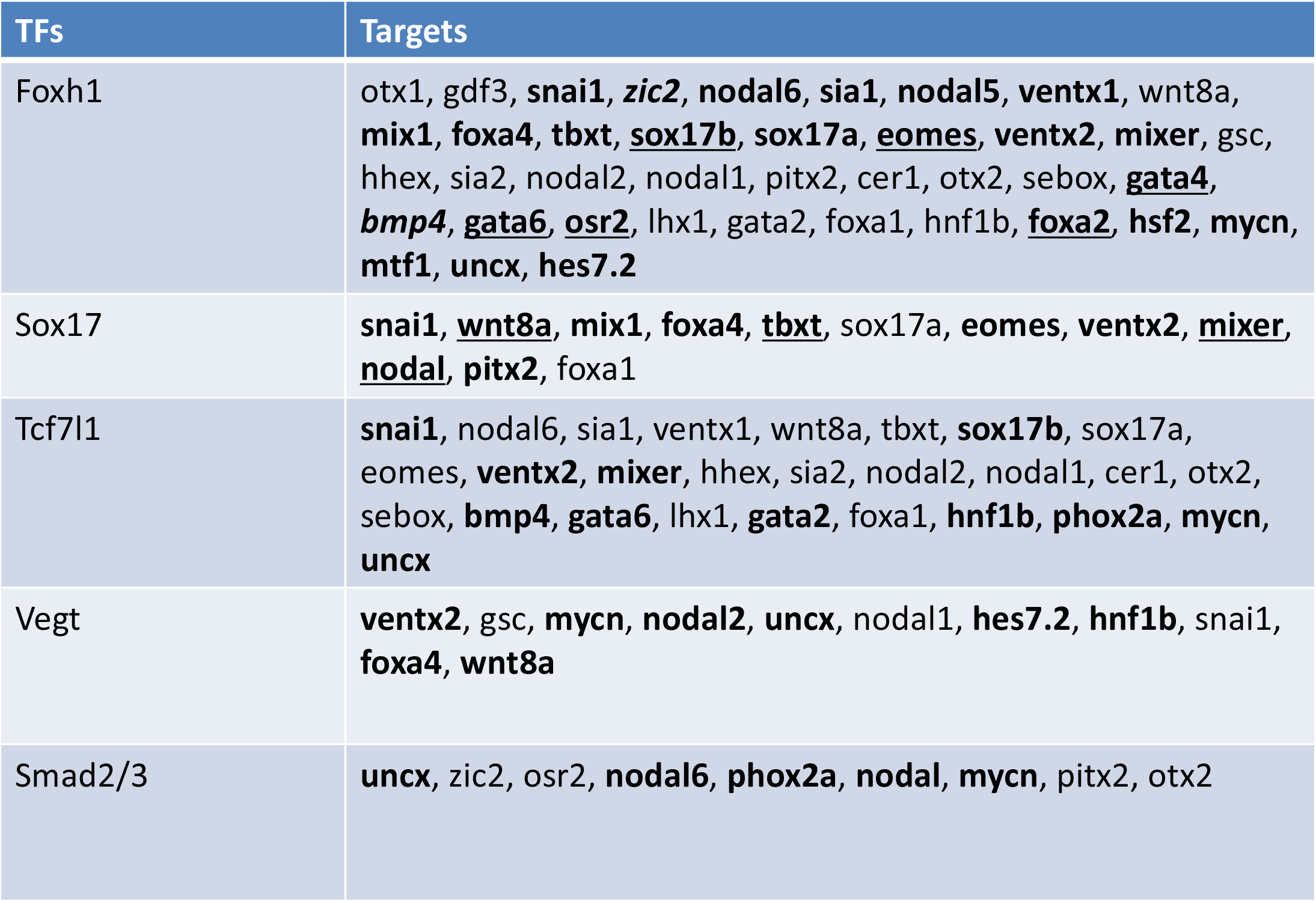
New and known core mesendodermal TF targets. List of targets in the core mesendodermal network for the TFs: Foxh1, Sox17, and Tcf7l1. Bolded entries are new to this analysis. Underlined entries were successfully validated. Italicized entries were inconclusively validated.

In order to examine the accuracy of this CRM prediction in vivo, 12 newly identified, putative Foxh1 and Sox17 CRM targets were cloned upstream of a minimal *gsc* promoter driving a luciferase reporter gene. We tested for luciferase activity in the presence and absence of endogenous Foxh1 and Sox17. Specifically, either control, Foxh1 or Sox17 MO was injected in the mesendoderm region of embryos and luciferase activities were compared (see STAR Methods). Tcf CRMs were not tested because 4 Tcf family TFs are expressed maternally and knocking down the expression of all of these by antisense MO has been difficult. Each microinjected CRM reporter showed expression in the mesendoderm (Figure 6B-C, Table S9, Figure S5B), which suggested that each probed region is a functional enhancer. Additionally, with the exception of two inconclusive reporters (from the *zic2* and *bmp4* genes, which were inconsistent between replicates), each reporter’s activity in mesendodermal cells was dependent to varying degrees on the presence of Foxh1 or Sox17. All reporter gene activities were downregulated in the absence of Foxh1 or Sox17, except *nodal*, suggesting that Foxh1 and Sox17 predominantly function as an activator for the genes belonging to metacluster R38 and R16. The *nodal* gene belongs to metacluster R82, and at present it is unclear whether the genes belonging to R82 are negatively regulated by Sox17 or whether this is due to an isolated action of this *nodal* CRM. Taken together, we conclude that the Linked SOM method of regulatory prediction combined with our new filtering methods shows a high-fidelity rate (10 confirmed cases out of 12 tested), while producing significantly more TF-CRM connections than previous methods.

## Discussion

Elucidating transcriptional GRNs is critical to understanding the molecular mechanisms of cell differentiation in developmental and disease systems. In recent years, significant efforts have been made to incorporate multi-omic datasets into GRN models, but this has been a difficult endeavor (Hu et al., 2020). In this work, in addition to a number of publicly available *Xenopus* genomic datasets, we generated additional RNA-seq and ChIP-seq data. We then combined 2 different SOM analyses, chromatin regulatory information (ChIP/ATAC-seq) and transcriptomic information (RNA-seq), to prioritize and identify key mesendodermal transcription factor targets. This integrated multi-omic approach was successful in accurately recapitulating cellular differentiation programs through network analysis. The generated GRN was validated both experimentally and statistically, to provide a highly-confident set of predictions of gene regulation controlling *Xenopus* mesendoderm development. These predicted connections not only included the known core mesendoderm networks from previous works (Charney et al, 2017b; Paraiso et al., 2020), but also provided a significant number of new connections that were previously unknown. Our analysis represents one of the most data-driven and integrative attempts to recapitulate the GRN of an *in vivo* developmental system.

### Novel network targets of key mesendermal transcription factors

Numerous genomic analyses of individual TFs have been used to understand early *Xenopus* development (Yoon et al., 2011; Gentsch et al., 2013; Chiu et al., 2014; Yasuoka et al., 2014; Nakamura et al., 2016; Kjolby and Harland, 2017; Charney et al., 2017b; Paraiso et al., 2019; Gentsch et al., 2019; Mukherjee et al., 2020; Afouda et al., 2020). In these experiments, combining a single, or a few, ChIP-seq dataset(s) and RNA-seq datasets in wild type and perturbed states have been used to identify direct transcriptional targets of TFs. A major limitation of this type of analysis is that target identification using a combination of ChIP peaks and large gene expression differences in MO loss of function analysis could miss small expression differences. These differences may be small due to compensatory effects from other TFs. By using an integrative approach which contextualizes a single TF ChIP-seq binding site with the binding of a multitude of regulatory proteins and correlating the binding with the expression of nearby genes, we identify 2,747 potential actionable targets of Foxh1 by leveraging multiple large datasets.

Of the 40 genes from the core mesendoderm network (Charney et al, 2017b), 34 had predicted functional Foxh1 motifs (Table 1), among which 14 genes were previously confirmed FoxH1 targets. While some genes such as *cer1, lhx1, otx2, sebox* and were previously shown to be regulated directly by Foxh1, *bmp4, gata4, gata6*, and *osr2* were never implicated as direct Foxh1 targets. Additionally, the metacluster of these genes, R16, also included 9 additional predicted targets such as *hoxd1* and *irx2*, which are critical to axis and pattern formation, respectively. At present, their roles in early mesendoderm formation are currently unknown. Another interesting metacluster is R38, of which only one of the potential Foxh1 targets had previous evidence, *wnt8a*. It turned out that the majority of core mesendodermal genes in R38 (except *tbxt*) including *sox17a* and *b*, which is active in a different region from *wnt8a*, were found to be similarly targeted by Foxh1. Comparing the temporal profiles of R16 and R38 in Figure 2a shows that these clusters have very similar temporal profiles, except genes in R38 being expressed at a significantly higher level than those in R16. This suggests that while Foxh1 regulates the expression of these genes, underlying mechanisms regulating these two metaclusters are different.

The predicted mesendoderm network also indicated that most of the genes in R38 were regulated by Sox17, whereas none in R16 were predicted. Genes in R38 also maintained a higher average gene expression level than those in R16. One speculation is that this difference in gene expression level is due to the positive feedback loop of Sox17 (Sinner et al., 2004; Howard et al., 2007) pulling each of these genes in lockstep with its expression. We tested the model using reporter genes driven by the CRMs of *mixer, tbxt* and *wnt8a* and validated that the output is regulated by both Foxh1 and Sox17 TF input *in vivo* (Figure S5A, Figure 5C). Additionally, the stage 10 expression of genes in metacluster R1 (in particular *snai1*) peak at nearly the same time point as R38. This is the only maternally and zygotically expressed metacluster with this peak and was the only one predicted to also be regulated by Sox17. Based on the current validation experiments, we also conclude that many of the newly predicted interactions between TFs and cis-regulatory modules are likely to have relevant function *in vivo*.

### Enhanceosomes, cooperativity and antagonism

While the main focus of this work was to elucidate the important cis-regulatory modules for gene regulation, an important component of the Linked SOM analysis, the ATAC/ChIP-seq SOM, revealed interesting clustering of TF binding. Importantly, the output of this SOM has shown consistency with known TF-TF interactions such as that of endodermal maternal TFs (Paraiso et al., 2019), Spemann organizer genes (Yasuoka et al., 2014), and mesodermal T-box genes (Gentsch, et al., 2013). This unbiased multi-omic clustering approach renders support for the importance of these respective enhanceosomes, complexes of TFs around enhancers. In the future, chromatin clustering with additional data is likely to reveal other interesting enhanceosome biology relevant to development.

Enhanceosomes positively regulate gene expression. The Ep300 co-activator is a histone acetyltransferase and its interaction with CRMs is one of the frequently used genomic markers of enhancer regions (Heintzman et al., 2007). MEME analysis of Ep300 peaks reveals the enrichment of Sox and Fox TF binding motifs, indicating that Ep300 is recruited to DNA via Sox and/or Fox family TFs. Consistent with this observation, we find that early Ep300 binding clusters with Foxh1 (at Stage 9) and late Ep300 binding clusters with Foxa2 (at Stage 10). Interestingly, Ep300 did not cluster with Sox7 and Sox17, indicating that other Sox family TFs, such as Sox3, may be responsible for this recruitment. We also note that Smad2/3 binding, which is a sign of Foxh1 mediated Nodal signaling activity, had a very poor correlation with Ep300 (of the 7707 Smad2/3 CRMs, only 41 overlapped with a significant Ep300 ChIP signal). This suggests that Ep300 interaction is dynamic – It is initially recruited to the potential sites by maternally expressed Sox and Fox TFs and gradually replaced by other zygotic TFs as Foxa2.

Our ATAC/ChIP-seq SOM revealed surprisingly close hierarchical clustering of ChIP signals for TFs that have distinct spatial expression differences (Figure S3). One of three examples includes the cluster containing dorsally expressed regulator of the Nieuwkoop organizing center homeobox Sia1 (Lemaire et al., 1995) and the ventrally expressed homeobox Ventx2 (Schmidt et al., 1996). This is unexpected as these TFs are known to specify opposing cell types (dorsal vs. ventral) and known to be expressed in spatially distinct embryonic regions. One possibility is that these two homeobox TFs bind competitively to similar motifs, and recruit two distinct enhanceosomes to the same enhancers, depending on the cellular environment. For instance, Sia1 may activate a subset of genes through these enhancers, whereas in a different region of the embryo, Ventx2 may use these same enhancers to repress target genes via recruiting a different combination of co-factors. Alternatively, these enhancers could be similarly regulated in dorsal and ventral regions of the embryo by Sia1 or Ventx2, but other spatial-specific factors could change the topology of the chromatin to target two distinct sets of genes from the same enhancer. Secondly, we identified this same pattern in other dorsal-ventral pairs of TFs such as the signaling pathway transducers Ctnnb1 (β-catenin; Wnt signaling transducer) (Stevens el al., 2017; Heasman et al., 1994), Foxh1 (Nodal signaling co-factor) (Chen et al., 1996), and Smad1 (Bmp signaling transducer) (Graff et al., 1996). The first two are both important for establishing the dorsal domain of the embryo, while Smad1 helps establish ventral identity. Finally, we note a similar pattern for the TFs Sox17 (Hudson et al., 1997) and Tbxt (Smith et al., 1991), which are critical TFs in forming the endoderm and mesoderm, respectively. A further focused study on these competitive binding locations could help answer how cells dynamically regulate gene expression by sharing similar enhanseosome modules during gastrulation.

In conclusion, we show that Linked SOMs are capable of efficiently predicting biologically relevant transcription factor-enhancer interactions to understand gene regulatory mechanism in an archetypical developmental system. To do this, our approach used a multi-omic dataset to create a highly-accurate mechanistic GRN without converting our ChIP/ATAC-seq data into RNA-seq-like data. These results cemented the important role of endodermal TFs, such as Foxh1 and Sox17, in coordinating the expression of many important developmental genes. Our work provides a useful, new platform for the data integration of multi-omic datasets (ChIP-seq, ATAC-seq, RNA-seq, etc) to uncover transcription factor-enhancer interactions *in vivo* cell and developmental systems. While we have applied Linked SOM for bulk sequencing data, the approach is flexible and can easily integrate other datasets such as single cell sequencing datasets.

## Acknowledgements

This work was made possible, in part, through access to the Genomic High-Throughput Facility Shared Resource of the Cancer Center Support Grant (P30CA-062203) at the University of California, Irvine, and NIH shared instrumentation grants 1S10RR025496-01, 1S10OD010794-01, and S10OD021718-01. We thank Xenbase for genomic and community resources (http://www.xenbase.org/entry, RRID: SCR_003280), in addition to their bioinformatic assistance; Amphibian Research Center at Hiroshima University (JP19km0210085); and the University of California, Irvine High Performance Computing Cluster (https://hpc.oit.uci.edu/) for their valuable resources and helpful staff. We also thank Y. Honda and Y. Kirigaya for Sia, Mix and Vegt antibody and ChIP testing. This research was funded by the following grants awarded to K.W.Y.C.: NIH R01HD073179, R01GM126395 and National Science Foundation (NSF) 1755214. K.D.P. was a recipient of NIH T32-HD60555.

## Author Contributions

A.M., I.L.B., and K.W.Y.C. conceived the project and developed the overall methods. K.D.P., J.J.Z., I.L.B., M.B.F., R.M.C., J.S.C., Y.Y., N.S., A.R.B. and M.W. performed bench experiments. K.D.P performed curation and preliminary analysis of the genomic datasets. A.M. and C.J. developed and performed the SOM analysis. A.M., I.L.B., and K.W.Y.C. supervised the research and co-wrote the manuscript with C.D.J. and K.D.P. Y.Y., G.J.C.V., A.M. and M.T. contributed to discussion and editing of the manuscript.

## Declaration of Interests

The authors declare no competing interests.

## STAR METHODS

### LEAD CONTACT AND MATERIALS AVAILABILITY

Further information and requests for resources and reagents should be directed to and will be fulfilled by the Lead Contact, Ken W.Y. Cho.

### EXPERIMENTAL MODEL AND SUBJECT DETAILS

Wild type *Xenopus tropicalis* males and females were either obtained from NASCO (University of Virginia) or raised in the laboratory and were maintained in accordance to the University of California, Irvine Institutional Animal Care Use Committee (IACUC). *X. tropicalis* females were injected with 10 units of Chorulon HCG 1-3 nights prior to use, and were injected with 100 units of Chorulon HCG the morning of use. Eggs were collected into a glass dish coated with 0.1% BSA in 1/9x MMR. Sperm suspension obtained from sacrificed males was used to *in vitro* fertilize the eggs. Ten minutes after fertilization, the embryos were dejellied with 3% cysteine in 1/9x MMR, pH 7.8 and are then ready for further manipulation.

### METHOD DETAILS

#### ChIP-seq and ATAC-seq

Majority of ChIP-seq datasets were obtained from NCBI’s Gene Expression Omnibus (see Key Resources Table). For newly generated datasets, ChIP-seq was performed as previously described (Chiu et al., 2014) at the appropriate developmental stage. The antibodies and conditions for these datasets:

- 30 ug of published VegT antibody (Sudou et al., 2012) per 2000-3000 embryos
- 30 ug of published Mix1 antibody (Sudou et al., 2012) per 2000-3000 embryos
- 30 ug of published Sia1 antibody (Sudou et al., 2012) per 2000-3000 embryos
- 4ug of Sox7 rabbit polyclonal peptide antibody (Genscript) per 100 embryos; the peptide antibody was designed against a region in the Sox7 transactivation domain in the C-terminus with the sequence QVSQASDIQPSETS
- 3.5 ug of Ventx2 rabbit polyclonal antibody per 100 embryos; the antibody was generated by Covance, Inc., using a GST fusion to Ventx2.2 amino acids 2-153, upstream of the homeodomain.
- 2.5 ug of Smad1/5/8 antibody (Santa Cruz Biotechnology sc-6031x) per 100 embryos

Libraries were generated using NEXTflex ChIP-seq (Bioo Scientific, Cat# NOVA-5143-01) kit, quality tested using an Agilent Bioanalyzer 2100, quantified using KAPA qPCR, and sequenced using Illumina sequencers at the UC Irvine Genomics High Throughput Facility.

ATAC-seq was available from BioRxiv: doi.org/10.1101/2020.02.26.966168 (Bright et al.,2020).

#### Gene knockdown and RNA-seq

Published RNA-seq datasets for different embryonic tissues and experimental conditions were obtained from NCBI’s Gene Expression Omnibus (see Key Resources Table). For the MO experiments, 2ng/embryo of *ctnnb1* MO (Mukherjee et al., 2020), 20 ng/embryo *foxh1* MO (Chiu et al., 2014; Charney et al, 2017), 10 ng/embryo each of two *sox17* MOs (targeting sox17a and sox17b1/2) or 4 ng/embryo *tcf7l1* MO (Liu et al., 2005) were used. For the knockdown of receptor-mediated Smad2/3 phosphorylation, embryos were treated with SB4315422 at 100uM as previously described (Chiu et al., 2014; Charney et al., 2017). For each condition, embryos were harvested at the appropriate developmental stage adhering to the *Xenopus* developmental table (Nieuwkoop and Faber, 1958). RNA samples were collected from embryos using the acid guanidium isothiocyanate method (Chomczynski and Sacchi, 1987). RNA-seq libraries were generated using Smart-seq2 cDNA synthesis followed by tagmentation (Picelli et al., 2014), quality tested using an Agilent Bioanalyzer 2100, quantified using KAPA qPCR, and sequenced using Illumina sequencers at the UC Irvine Genomics High Throughput Facility.

#### Construction of Luciferase Reporter Assay Plasmids

Minimal *gsc* promoter (−104*gsc*) is PCR amplified from −104gsc/Luc (Watabe et al. 1995) and cloned into pGL3 basic vector (Promega) at HindIII and NcoI restriction digestion sites.

Candidate foxh1-bound CRMs are PCR amplified from *Xenopus* tropicalis genomic DNA (primers are listed in Table S10) and cloned in above vector at BglII and HindIII restriction digestion sites.

#### Luciferase Assay for Validation of Predicted Foxh1-bound CRMs

To examine the activity of each Foxh1-bound CRM, 20pg of luciferase reporter construct and 2pg of pRL-SV40 (Promega) are co-injected vegetally into a 1-cell stage embryo. Luciferase reporter construct without a CRM is served as a negative control. Injected embryos are harvested at stage 10.5 (early gastrula) by homogenizing 10 embryos in 100ul of 5X passive lysis buffer (Promega). 30ul of clear lysate cleared of cellular debris are used to assay for luciferase activities according to the manufacturer instruction of Dual-Luciferase Reporter Assay System (Promega).

### QUANTIFICATION AND STATISTICAL ANALYSIS

#### Chromatin Dataset Analysis

ATAC-seq and ChIP-seq reads were aligned to the *X. tropicalis* genome v 9.0 (Mitros et al., 2019) obtained from Xenbase (Karimi et al., 2018) using Bowtie 2 v2.2.7 (Langmed and Salzberg et al., 2012). ATAC-seq and ChIP-seq datasets were peak called relative to their appropriate input DNA controls using MACS2 v.2.0.10 (Zhang et al., 2008) with default options.

#### Chromatin segmentation and DNA-SOM analysis

The *Xenopus tropicalis* v9 reference genome was partitioned using the **partition** tool of SOMatic (Jansen et al., 2019) using the MACS2 peak files with a minimum partition size of 200 bp. Then, a RPKM matrix was calculated using the **regionCounts** tool from SOMatic.

The DNA SOM was built using the **buildSite** tool from SOMatic, using a size of 40×60, 100 epochs, 100 trials. SOMatic found 88 metaclusters had the best AIC score using 100 trials. GO term enrichments were found using the XenMine gene ontology tool (Reid et al., 2017).

#### RNA-seq Dataset Analysis

RNA-seq reads were aligned to the *X. tropicalis* genome v 9.0 (Mitros et al., 2019) obtained from Xenbase (Karimi et al., 2018) using RSEM v 1.2.12 (Li and Dewey, 2011) and Bowtie 2 v2.2.7 (Langmed and Salzberg et al., 2012) to generate gene expression in transcripts per million.

#### RNA-SOM analysis

The RNA SOM was built using the buildSite tool from SOMatic, using a size of 60×90, 100 epochs, 100 trials. SOMatic found 84 metaclusters had the best AIC score using 100 trials. Various SOMatic tools were used to create all of the heatmaps, including the statistical enrichment graph, and GO term enrichments were found using the Xenbase GO term tool.

#### Linking of DNA- and RNA-SOM and Network analysis

The **Link** tool in SOMatic was used to convolve the 2 SOMs’ metaclusters, using the nearest gene option and limiting the search area to 1Mb. A specific *Xenopus* option (-Xeno) was used because the Xenbase gtf file is a non-standard format.

For the initial ChIP/ATAC-seq SOM, the regions, including repeat regions, in each metacluster were scanned for motifs using the HOCOMOCOv11 human motif database (Kulakovskiy et al., 2017) with FIMO v4.12.0 using a q-value threshold of 0.1. For the further network analysis, each linked metacluster (LM) was scanned with FIMO v4.12.0 (Grant et al., 2011) using a q-value threshold of .1 using motifs calculated from the *Xenopus* ChIP data. The background for both analyses was calculated using the entire *Xenopus tropicalis* v9 reference genome. For each of the 12 calculated TF motifs, the percentage of regions in each LM with that motif was calculated and used to perform one-tailed z-score enrichment with a q-value of 0.05. These significant TF motif locations were mapped to the linked gene.

#### Gene enrichment analysis for unchanging genes throughout time-course

We used DESeq2 v3.11 (Love et al., 2014) to find significantly unchanging genes by using the altHypothesis=“lessAbs” option (qvalue < .05).

### DATA AND CODE AVAILABILITY

The github for SOMatic can be found here: https://github.com/csjansen/SOMatic. Raw and processed RNA-seq and ChIP-seq datasets generated for this study are available at NCBI Gene Expression Omnibus using the accession GEO: TBD.

## Supplemental information titles and legends

**Figure S1. *SOM slices reveal overall structure of the input data*. (A)** An RNA SOM slice corresponding to wildtype (stage 10.5). The SOM unit location of various important genes in mesendodermal development are noted. **(B)** RNA SOM difference slices corresponding to the fold change between MO and control experiments. The RNA metacluster divisions are overlaid over the slices. **(C)** DNA SOM slices corresponding to Foxh1, Vegt, and Otx1 ChIPs at stage 8 and one corresponding to their average. The DNA metacluster divisions are overlaid over the slices. **(D)** DNA SOM slices corresponding to Foxh1 and Ep300 ChIPs at stage 9 and their average, followed by DNA SOM slices corresponding to Smad2/3, Foxh1, and Gsc ChIPs at stage 10.5. The DNA metacluster divisions are overlaid over the slices. **(E)** DNA SOM slices corresponding to Sox17 ChIP at stage 10.5 and Tbxt ChIP at stage 12. The DNA metacluster divisions are overlaid over the slices. **(F)** DNA SOM slices corresponding to Ctnnb1, Foxh1, and Smad1 ChIPs at stage 10.5. The DNA metacluster divisions are overlaid over the slices.

**Figure S2. *Distribution of TF ChIP peak summits in final partitioning*.** The number of unique TF peak summits within each partition. 46% of partitions contain at least one TF peak summit.

**Figure S3. *Full DNA metacluster heatmap captures known co-regulatory interactions*.** The full set of eigenprofiles revealed that several experiments had very similar results on the collected genomic region clusters. Some of these are known co-regulatory interactions in *Xenopus* or vertebrates in general

**Figure S4. *Motif analysis on linked spatial RNA and DNA SOM metaclusters finds TFs with spatially specific regulation*. (A)** Venn diagrams showing the motif overlap between spatial metaclusters with similar temporal gene expression profiles. **(B)** The overlaps between motifs found uniquely on each side of the embryo in (A).

**Figure S5. *Final network luciferase validation experiments*. (A)** Motif filtering strategy of network analysis. After using FIMO to find motifs and Motif Enrichment to filter out motifs that are dispersed evenly across the linked metaclusters, the network connections were further filtered by a joint Ep300 and TF ChIP signal separately for each of the TFs tested. Then, we selected RNA metaclusters that contained TFs known to be important for mesendodermal development. Finally, we connected TFs to genes from the core mesendodermal network and known TFs to make the final network. **(B)** A genome browser view of the predicted and tested Foxh1 target element near *gata6* with Foxh1, Sox17, Ctnnb1 (β-catenin), and Ep300 ChIP-seq signals for stages 8 - 12.

## Notes

### Competing Interest Statement

The authors have declared no competing interest.

## References

Afouda, B.A., Ciau-Uitz, A., and Patient, R. (2005). GATA4, 5 and 6 mediate TGFbeta maintenance of endodermal gene expression in *Xenopus* embryos. Development 132, 763–774.

Afouda, B.A., Nakamura, Y., Shaw, S., Charney, R.M., Paraiso, K.D., Blitz, I.L., Cho, K.W.Y., and Hoppler, S. (2020). Foxh1/Nodal Defines Context-Specific Direct Maternal Wnt/β-Catenin Target Gene Regulation in Early Development. iScience 23, 101314.

Ault, K.T., Dirksen, M.L., and Jamrich, M. (1996). A novel homeobox gene PV.1 mediates induction of ventral mesoderm in *Xenopus* embryos. Proc Natl Acad Sci USA 93, 6415–6420.

Bardet, A.F., Steinmann, J., Bafna, S., Knoblich, J.A., Zeitlinger, J., and Stark, A. (2013). Identification of transcription factor binding sites from ChIP-seq data at high resolution. Bioinformatics 29, 2705–2713.

Bergsland, M., Werme, M., Malewicz, M., Perlmann, T., and Muhr, J. (2006). The establishment of neuronal properties is controlled by Sox4 and Sox11.Genes & Development 20, 3475–3486.

Blitz, I.L., Paraiso, K.D., Patrushev, I., Chiu, W.T.Y., Cho, K.W.Y., and Gilchrist, M.J. (2017). A catalog of *Xenopus tropicalis* transcription factors and their regional expression in the early gastrula stage embryo. Dev Biol 426, 409–417.

Bright, A.R., Genesen, S. van, Li, Q., Heeringen, S.J. van, Grasso, A., and Veenstra, G.J.C. (2020). Combinatorial action of transcription factors in open chromatin contributes to early cellular heterogeneity and organizer mesendoderm specification. BioRxiv DOI: https://doi.org/10.1101/2020.02.26.966168

Buenrostro, J.D., Giresi, P.G., Zaba, L.C., Chang, H.Y., and Greenleaf, W.J. (2013). Transposition of native chromatin for fast and sensitive epigenomic profiling of open chromatin, DNA-binding proteins and nucleosome position. Nat Methods 10, 1213–1218.

Cao, R., Wang, L., Wang, H., Xia, L., Erdjument-Bromage, H., Tempst, P., Jones, R.S., and Zhang, Y. (2002). Role of histone H3 lysine 27 methylation in Polycomb-group silencing. Science 298, 1039–1043.

Charney, R.M., Paraiso, K.D., Blitz, I.L., and Cho, K.W.Y. (2017a). A gene regulatory program controlling early *Xenopus* mesendoderm formation: Network conservation and motifs. Semin Cell Dev Biol 66, 12–24.

Charney, R.M., Forouzmand, E., Cho, J.S., Cheung, J., Paraiso, K.D., Yasuoka, Y., Takahashi, S., Taira, M., Blitz, I.L., Xie, X., et al. (2017b). Foxh1 Occupies cis-Regulatory Modules Prior to Dynamic Transcription Factor Interactions Controlling the Mesendoderm Gene Program. Dev Cell 40, 595–607.e4.

Chen, C., Jin, J., Lee, G.A., Silva, E., and Donoghue, M. (2016). Cross-species functional analyses reveal shared and separate roles for Sox11 in frog primary neurogenesis and mouse cortical neuronal differentiation. Biology Open 5, 409–417.

Chen, X., Rubock, M.J., and Whitman, M. (1996). A transcriptional partner for MAD proteins in TGF-beta signalling. Nature 383, 691–696.

Cheng, Y., Ma, Z., Kim, B.H., Wu, W., Cayting, P., Boyle, A.P., Sundaram, V., Xing, X., Dogan, N., Li, J., et al. (2014). Principles of regulatory information conservation between mouse and human. Nature 515, 371–375.

Chiu, W.T., Charney, L.R., Blitz, I.L., Fish, M.B., Li, Y., Biesinger, J., Xie, X., and Cho, K.W. (2014). Genome-wide view of TGFβ/Foxh1 regulation of the early mesendoderm program. Development 141, 4537–4547.

Cho, K.W., Blumberg, B., Steinbeisser, H., and De, R.E.M. (1991). Molecular nature of Spemann’s organizer: the role of the *Xenopus* homeobox gene goosecoid. Cell 67, 1111–1120.

Chomczynski, P., and Sacchi, N. (1987). Single-step method of RNA isolation by acid guanidinium thiocyanate-phenol-chloroform extraction. Analytical Biochemistry 162, 156–159.

Conlon, F.L., Fairclough, L., Price, B.M., Casey, E.S., and Smith, J.C. (2001). Determinants of T box protein specificity. Development 128, 3749–3758.

The ENCODE Project Consortium (2012). An integrated encyclopedia of DNA elements in the human genome. Nature 489, 57–74.

Creyghton, M.P., Cheng, A.W., Welstead, G.G., Kooistra, T., Carey, B.W., Steine, E.J., Hanna, J., Lodato, M.A., Frampton, G.M., Sharp, P.A., et al. (2010). Histone H3K27ac separates active from poised enhancers and predicts developmental state. Proc Natl Acad Sci USA 107, 21931–21936.

Davidson, E.H. (2010). Emerging properties of animal gene regulatory networks. Nature 468, 911–920.

Delgado, F.M., and Gómez-Vela, F. (2019). Computational methods for Gene Regulatory Networks reconstruction and analysis: A review. Artificial Intelligence in Medicine 95, 133–145.

Edgar, R., Mazor, Y., Rinon, A., Blumenthal, J., Golan, Y., Buzhor, E., Livnat, I., Ben-Ari, S., Lieder, I., Shitrit, A., et al. (2013). LifeMap Discovery: The Embryonic Development Stem Cells, and Regenerative Medicine Research Portal. PLoS ONE 8, e66629.

Ernst, J., and Kellis, M. (2012). ChromHMM: automating chromatin-state discovery and characterization. Nat Methods 9, 215–216.

Gurdon, J.B., Elsdale, T.R., and Fischberg, M. (1958). Sexually mature individuals of *Xenopus laevis* from the transplantation of single somatic nuclei. Nature 182, 64–65.

Gentsch, G.E., Owens, N.D., Martin, S.R., Piccinelli, P., Faial, T., Trotter, M.W., Gilchrist, M.J., and Smith, J.C. (2013). In vivo T-box transcription factor profiling reveals joint regulation of embryonic neuromesodermal bipotency. Cell Rep 4, 1185–1196.

Gentsch, G.E., Spruce, T., Owens, N.D.L., and Smith, J.C. (2019). Maternal pluripotency factors initiate extensive chromatin remodelling to predefine first response to inductive signals. Nat Commun 10, 4269.

Graff, J.M., Bansal, A., and Melton, D.A. (1996). *Xenopus* Mad proteins transduce distinct subsets of signals for the TGF beta superfamily. Cell 85, 479–487.

Grant, C.E., Bailey, T.L., and Noble, W.S. (2011). FIMO: scanning for occurrences of a given motif. Bioinformatics 27, 1017–1018.

Heasman, J., Crawford, A., Goldstone, K., Garner-Hamrick, P., Gumbiner, B., McCrea, P., Kintner, C., Noro, C.Y., and Wylie, C. (1994). Overexpression of cadherins and underexpression of beta-catenin inhibit dorsal mesoderm induction in early *Xenopus* embryos. Cell 79, 791–803.

Heintzman, N.D., Stuart, R.K., Hon, G., Fu, Y., Ching, C.W., Hawkins, R.D., Barrera, L.O., Van, C.S., Qu, C., Ching, K.A., et al. (2007). Distinct and predictive chromatin signatures of transcriptional promoters and enhancers in the human genome. Nat Genet 39, 311–318.

Hontelez, S., van, K.I., Georgiou, G., van, H.S.J., Bogdanovic, O., Lister, R., and Veenstra, G.J.C. (2015). Embryonic transcription is controlled by maternally defined chromatin state. Nat Commun 6, 10148.

Howard, L., Rex, M., Clements, D., and Woodland, H.R. (2007). Regulation of the *Xenopus* Xsox17alpha(1) promoter by co-operating VegT and Sox17 sites. Dev Biol 310, 402–415.

Hu, X., Hu, Y., Wu, F., Leung, R.W.T., and Qin, J. (2020). Integration of single-cell multi-omics for gene regulatory network inference. Computational and Structural Biotechnology Journal 18, 1925–1938.

Hudson, C., Clements, D., Friday, R.V., Stott, D., and Woodland, H.R. (1997). Xsox17alpha and -beta mediate endoderm formation in *Xenopus*. Cell 91, 397–405.

Izzi, L., Silvestri, C., von Both, I., Labbé, E., Zakin, L., Wrana, J.L., Attisano, L. (2007). Foxh1 recruits Gsc to negatively regulate Mixl1 expression during early mouse development. EMBO J. 26, 3132–43.

Jansen, C., Ramirez, R.N., El-Ali, N.C., Gomez-Cabrero, D., Tegner, J., Merkenschlager, M., Conesa, A., and Mortazavi, A. (2019). Building gene regulatory networks from scATAC-seq and scRNA-seq using Linked Self Organizing Maps. PLoS Comput Biol 15, e1006555.

Karimi, K., Fortriede, J.D., Lotay, V.S., Burns, K.A., Wang, D.Z., Fisher, M.E., Pells, T.J., James-Zorn, C., Wang, Y., Ponferrada, V.G., et al. (2017). Xenbase: a genomic epigenomic and transcriptomic model organism database. Nucleic Acids Research 46, D861–D868.

Kiang, M.Y., and Kumar, A. (2001). An Evaluation of Self-Organizing Map Networks as a Robust Alternative to Factor Analysis in Data Mining Applications. Information Systems Research 12, 177–194.

Kjolby, R.A.S., and Harland, R.M. (2017). Genome-wide identification of Wnt/β-catenin transcriptional targets during *Xenopus* gastrulation. Dev Biol 426, 165–175.

Kohonen, T. (2001). Self-Organizing Maps (Springer Berlin Heidelberg).

Koide, T., Hayata, T., and Cho, K.W. (2005). *Xenopus* as a model system to study transcriptional regulatory networks. Proc Natl Acad Sci USA 102, 4943–4948.

Kulakovskiy, I.V., Vorontsov, I.E., Yevshin, I.S., Sharipov, R.N., Fedorova, A.D., Rumynskiy, E.I., Medvedeva, Y.A., Magana-Mora, A., Bajic, V.B., Papatsenko, D.A., et al. (2018). HOCOMOCO: towards a complete collection of transcription factor binding models for human and mouse via large-scale ChIP-Seq analysis. Nucleic Acids Res 46, D252–D259.

Kvon, E.Z., Stampfel, G., Yáñez-Cuna, J.O., Dickson, B.J., and Stark, A. (2012). HOT regions function as patterned developmental enhancers and have a distinct cis-regulatory signature. Genes Dev 26, 908–913.

Lagna, G., Hata, A., Hemmati-Brivanlou, A., and Massagué, J. (1996). Partnership between DPC4 and SMAD proteins in TGF-beta signalling pathways. Nature 383, 832–836.

Langmead, B., and Salzberg, S.L. (2012). Fast gapped-read alignment with Bowtie 2. Nature Methods 9, 357–359.

Lemaire, P., Garrett, N., and Gurdon, J.B. (1995). Expression cloning of Siamois, a *Xenopus* homeobox gene expressed in dorsal-vegetal cells of blastulae and able to induce a complete secondary axis. Cell 81, 85–94.

Levine, M., and Davidson, E.H. (2005). Gene regulatory networks for development. Proc Natl Acad Sci USA 102, 4936–4942.

Li, B., and Dewey, C.N. (2011). RSEM: accurate transcript quantification from RNA-Seq data with or without a reference genome. BMC Bioinformatics 12.

Li, X.Y., MacArthur, S., Bourgon, R., Nix, D., Pollard, D.A., Iyer, V.N., Hechmer, A., Simirenko, L., Stapleton, M., Luengo, H.C.L., et al. (2008). Transcription factors bind thousands of active and inactive regions in the Drosophila blastoderm. PLoS Biol 6, e27.

Lin, L., Lee, V.M., Wang, Y., Lin, J.S., Sock, E., Wegner, M., and Lei, L. (2010). Sox11 regulates survival and axonal growth of embryonic sensory neurons. Developmental Dynamics 240, 52–64.

Liu, F., van, den B.O., Destrée, O., and Hoppler, S. (2005). Distinct roles for *Xenopus* Tcf/Lef genes in mediating specific responses to Wnt/beta-catenin signalling in mesoderm development. Development 132, 5375–5385.

Lolas, M., Valenzuela, P.D., Tjian, R., and Liu, Z. (2014). Charting Brachyury-mediated developmental pathways during early mouse embryogenesis. Proc Natl Acad Sci USA 111, 4478–4483.

Loose, M., and Patient, R. (2004). A genetic regulatory network for *Xenopus* mesendoderm formation. Dev Biol 271, 467–478.

Love, M.I., Huber, W., and Anders, S. (2014). Moderated estimation of fold change and dispersion for RNA-seq data with DESeq2. Genome Biol 15, 550.

Maeda, T., Chapman, D.L., and Stewart, A.F. (2002). Mammalian vestigial-like 2, a cofactor of TEF-1 and MEF2 transcription factors that promotes skeletal muscle differentiation. J Biol Chem 277, 48889–48898.

Mitros, T., Lyons, J.B., Session, A.M., Jenkins, J., Shu, S., Kwon, T., Lane, M., Ng, C., Grammer, T.C., Khokha, M.K., et al. (2019). A chromosome-scale genome assembly and dense genetic map for *Xenopus tropicalis*. Developmental Biology 452, 8–20.

Mochizuki, T., Karavanov, A.A., Curtiss, P.E., Ault, K.T., Sugimoto, N., Watabe, T., Shiokawa, K., Jamrich, M., Cho, K.W., Dawid, I.B., et al. (2000). Xlim-1 and LIM domain binding protein 1 cooperate with various transcription factors in the regulation of the goosecoid promoter. Dev Biol 224, 470–485.

Mortazavi, A., Pepke, S., Jansen, C., Marinov, G.K., Ernst, J., Kellis, M., Hardison, R.C., Myers, R.M., and Wold, B.J. (2013). Integrating and mining the chromatin landscape of cell-type specificity using self-organizing maps. Genome Res 23, 2136–2148.

Mukherjee, S., Chaturvedi, P., Rankin, S.A., Fish, M.B., Wlizla, M., Paraiso, K.D., MacDonald, M., Chen, X., Weirauch, M.T., Blitz, I.L., et al. (2020). Sox17 and β-catenin co-occupy Wnt-responsive enhancers to govern the endoderm gene regulatory network. Elife 9: e58029.

Nakamura, Y., de, P.A.E., Veenstra, G.J., and Hoppler, S. (2016). Tissue- and stage-specific Wnt target gene expression is controlled subsequent to β-catenin recruitment to cis-regulatory modules. Development 143, 1914–1925.

Nieuwkoop, P.D., and Faber, J. (1958). Normal Table of *Xenopus laevis* (Daudin). Copeia 1958, 65.

Owens, N. D. L., Blitz, I. L., Lane, M. A., Patrushev, I., Overton, J. D., Gilchrist, M. J., et al. (2016). Measuring Absolute RNA Copy Numbers at High Temporal Resolution Reveals Transcriptome Kinetics in Development. Cell Reports, 14, 632–647.

Paraiso, K.D., Blitz, I.L., Coley, M., Cheung, J., Sudou, N., Taira, M., and Cho, K.W.Y. (2019). Endodermal Maternal Transcription Factors Establish Super-Enhancers during Zygotic Genome Activation. Cell Rep 27, 2962–2977.e5.

Paraiso, K.D., Cho, J.S., Yong, J., and Cho, K.W.Y. (2020). Early *Xenopus* gene regulatory programs, chromatin states, and the role of maternal transcription factors. Curr Top Dev Biol 139, 35–60.

Partridge, E.C., Chhetri, S.B., Prokop, J.W., Ramaker, R.C., Jansen, C.S., Goh, S.T., Mackiewicz, M., Newberry, K.M., Brandsmeier, L.A., Meadows, S.K., et al. (2020). Occupancy maps of 208 chromatin-associated proteins in one human cell type. Nature 583, 720–728.

Picelli, S., Faridani, O.R., K Björklund, Åsa, Winberg, G., Sagasser, S., and Sandberg, R. (2014). Full-length RNA-seq from single cells using Smart-seq2.Nature Protocols9, 171–181.

Pliner, H.A., Shendure, J., and Trapnell, C. (2019). Supervised classification enables rapid annotation of cell atlases. Nat Methods 16, 983–986.

Reid, C.D., Karra, K., Chang, J., Piskol, R., Li, Q., Li, J.B., Cherry, J.M., and Baker, J.C. (2017). XenMine: A genomic interaction tool for the *Xenopus* community. Developmental Biology 426, 155–164.

Santos-Rosa, H., Schneider, R., Bannister, A.J., Sherriff, J., Bernstein, B.E., Emre, N.C., Schreiber, S.L., Mellor, J., and Kouzarides, T. (2002). Active genes are tri-methylated at K4 of histone H3. Nature 419, 407–411.

Schmidt, J.E., von, D.G., and Kimelman, D. (1996). Regulation of dorsal-ventral patterning: the ventralizing effects of the novel *Xenopus* homeobox gene Vox. Development 122, 1711–1721.

Schotta, G., Lachner, M., Sarma, K., Ebert, A., Sengupta, R., Reuter, G., Reinberg, D., Jenuwein, T. (2004). A silencing pathway to induce H3-K9 and H4-K20 trimethylation at constitutive heterochromatin. Genes Dev. 18, 1251–1262.

Sebé-Pedrós, A., Ariza-Cosano, A., Weirauch, M.T., Leininger, S., Yang, A., Torruella, G., Adamski, M., Adamska, M., Hughes, T.R., Gómez-Skarmeta, J.L., et al. (2013). Early evolution of the T-box transcription factor family. Proc Natl Acad Sci USA 110, 16050–16055.

Sethi, A., Gu, M., Gumusgoz, E., Chan, L., Yan, K.-K., Rozowsky, J., Barozzi, I., Afzal, V., Akiyama, J.A., Plajzer-Frick, I., et al. (2020). Supervised enhancer prediction with epigenetic pattern recognition and targeted validation. Nature Methods 17, 807–814.

Sinner, D., Rankin, S., Lee, M., and Zorn, A.M. (2004). Sox17 and beta-catenin cooperate to regulate the transcription of endodermal genes. Development 131, 3069–3080.

Smith, J.C., Price, B.M., Green, J.B., Weigel, D., and Herrmann, B.G. (1991). Expression of a Xenopus homolog of Brachyury (T) is an immediate-early response to mesoderm induction. Cell 67, 79–87.

Stevens, M.L., Chaturvedi, P., Rankin, S.A., Macdonald, M., Jagannathan, S., Yukawa, M., Barski, A., and Zorn, A.M. (2017). Genomic integration of Wnt/β-catenin and BMP/Smad1 signaling coordinates foregut and hindgut transcriptional programs. Development 144, 1283–1295.

Stuart, T., Butler, A., Hoffman, P., Hafemeister, C., Papalexi, E., Mauck, W.M. 3rd, Hao, Y., Stoeckius, M., Smibert, P., and Satija, R. (2019). Comprehensive Integration of Single-Cell Data. Cell 177, 1888–1902.e21.

Sudou, N., Yamamoto, S., Ogino, H., and Taira, M. (2012). Dynamic in vivo binding of transcription factors to cis-regulatory modules of cer and gsc in the stepwise formation of the Spemann-Mangold organizer. Development 139, 1651–1661.

Takahashi, K., and Yamanaka, S. (2006). Induction of pluripotent stem cells from mouse embryonic and adult fibroblast cultures by defined factors. Cell 126, 663–676.

Veenstra, G.J., Beumer, T.L., Peterson-Maduro, J., Stegeman, B.I., Karg, H.A., van, der V.P.C., and Destrée, O.H. (1995). Dynamic and differential Oct-1 expression during early *Xenopus* embryogenesis: persistence of Oct-1 protein following down-regulation of the RNA. Mech Dev 50, 103–117.

Watabe, T., Kim, S., Candia, A., Rothbächer, U., Hashimoto, C., Inoue, K., and Cho, K.W. (1995). Molecular mechanisms of Spemann’s organizer formation: conserved growth factor synergy between *Xenopus* and mouse. Genes Dev 9, 3038–3050.

Welch, J.D., Kozareva, V., Ferreira, A., Vanderburg, C., Martin, C., and Macosko, E.Z. (2019). Single-Cell Multi-omic Integration Compares and Contrasts Features of Brain Cell Identity. Cell 177, 1873–1887.e17.

Wills, A.E., and Baker, J.C. (2015). E2a is necessary for Smad2/3-dependent transcription and the direct repression of lefty during gastrulation. Dev Cell 32, 345–357.

Xiang, G., Keller, C.A., Heuston, E., Giardine, B.M., An, L., Wixom, A.Q., Miller, A., Cockburn, A., Sauria, M.E.G., Weaver, K., et al. (2020). An integrative view of the regulatory and transcriptional landscapes in mouse hematopoiesis. Genome Research 30, 472–484.

Yasuoka, Y., Suzuki, Y., Takahashi, S., Someya, H., Sudou, N., Haramoto, Y., Cho, K.W., Asashima, M., Sugano, S., and Taira, M. (2014). Occupancy of tissue-specific cis-regulatory modules by Otx2 and TLE/Groucho for embryonic head specification. Nat Commun 5, 4322.

Yoon, S.J., Wills, A.E., Chuong, E., Gupta, R., and Baker, J.C. (2011). HEB and E2A function as SMAD/FOXH1 cofactors. Genes Dev 25, 1654–1661.

Yue, F., Cheng, Y., Breschi, A., Vierstra, J., Wu, W., Ryba, T., Sandstrom, R., Ma, Z., Davis, C., Pope, B.D., et al. (2014). A comparative encyclopedia of DNA elements in the mouse genome. Nature 515, 355–364.

Zhang, C., Basta, T., Fawcett, S.R., and Klymkowsky, M.W. (2005). SOX7 is an immediate-early target of VegT and regulates Nodal-related gene expression in *Xenopus*. Dev Biol 278, 526–541.

Zhang, Y., Liu, T., Meyer, C.A., Eeckhoute, J., Johnson, D.S., Bernstein, B.E., Nusbaum, C., Myers, R.M., Brown, M., Li, W., et al. (2008). Model-based analysis of ChIP-Seq (MACS). Genome Biol 9, R137.

Zorn, A.M., and Wells, J.M. (2009). Vertebrate endoderm development and organ formation. Annu Rev Cell Dev Biol 25, 221–251.

