## Supplementary material for "Uncovering the mesendoderm gene regulatory network through multi-omic data integration": Suppl figures

### Xenopus figures

Supp Figures

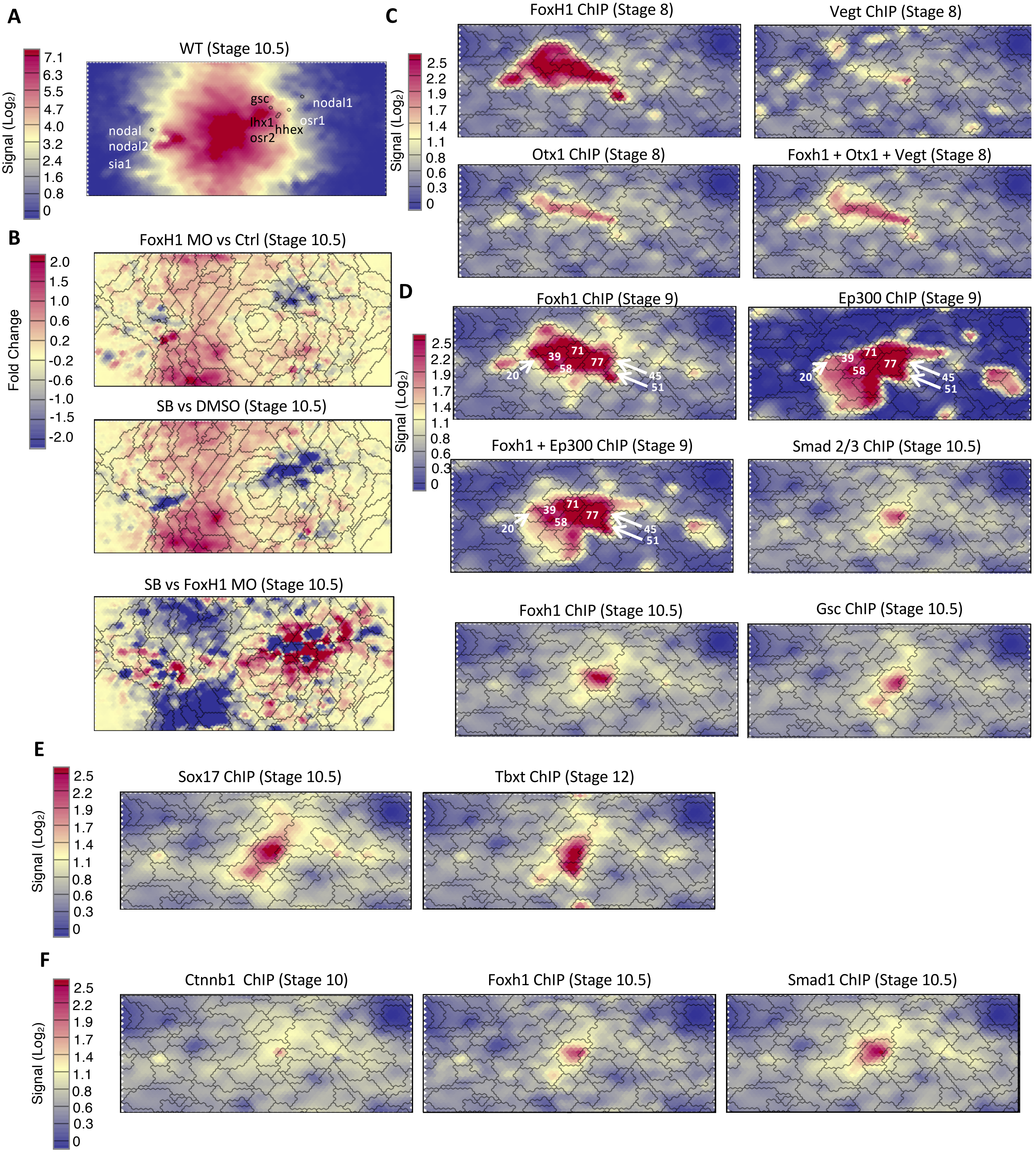


Suppl Figure 1

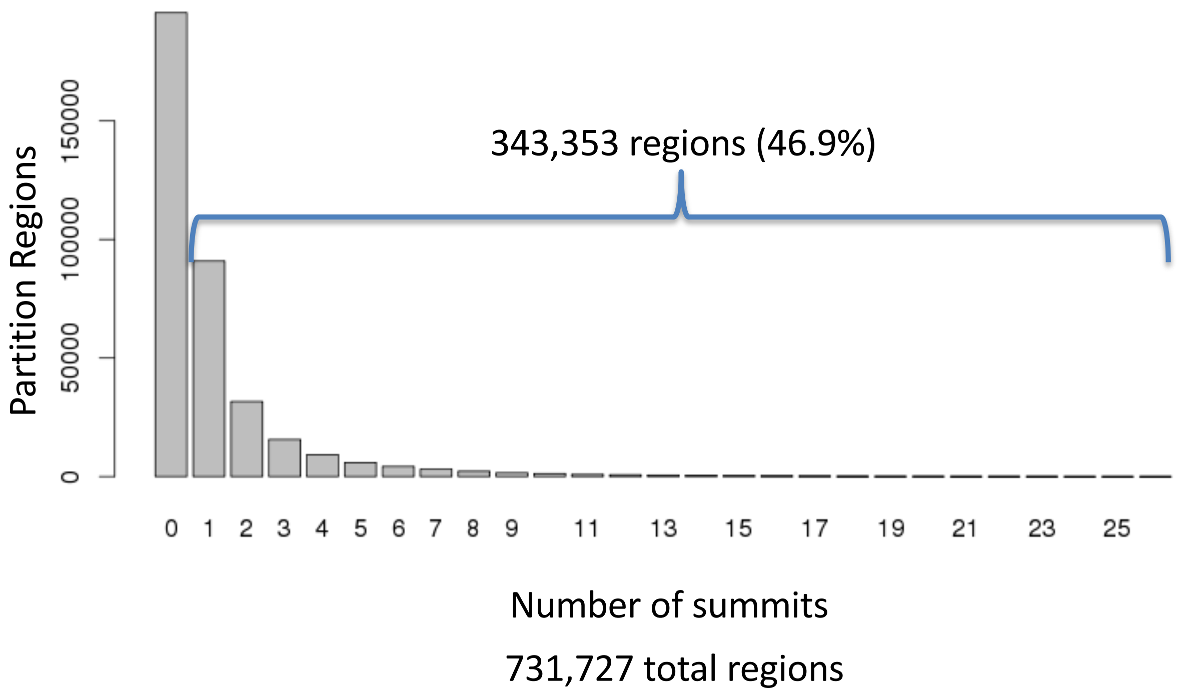


Suppl Figure 2

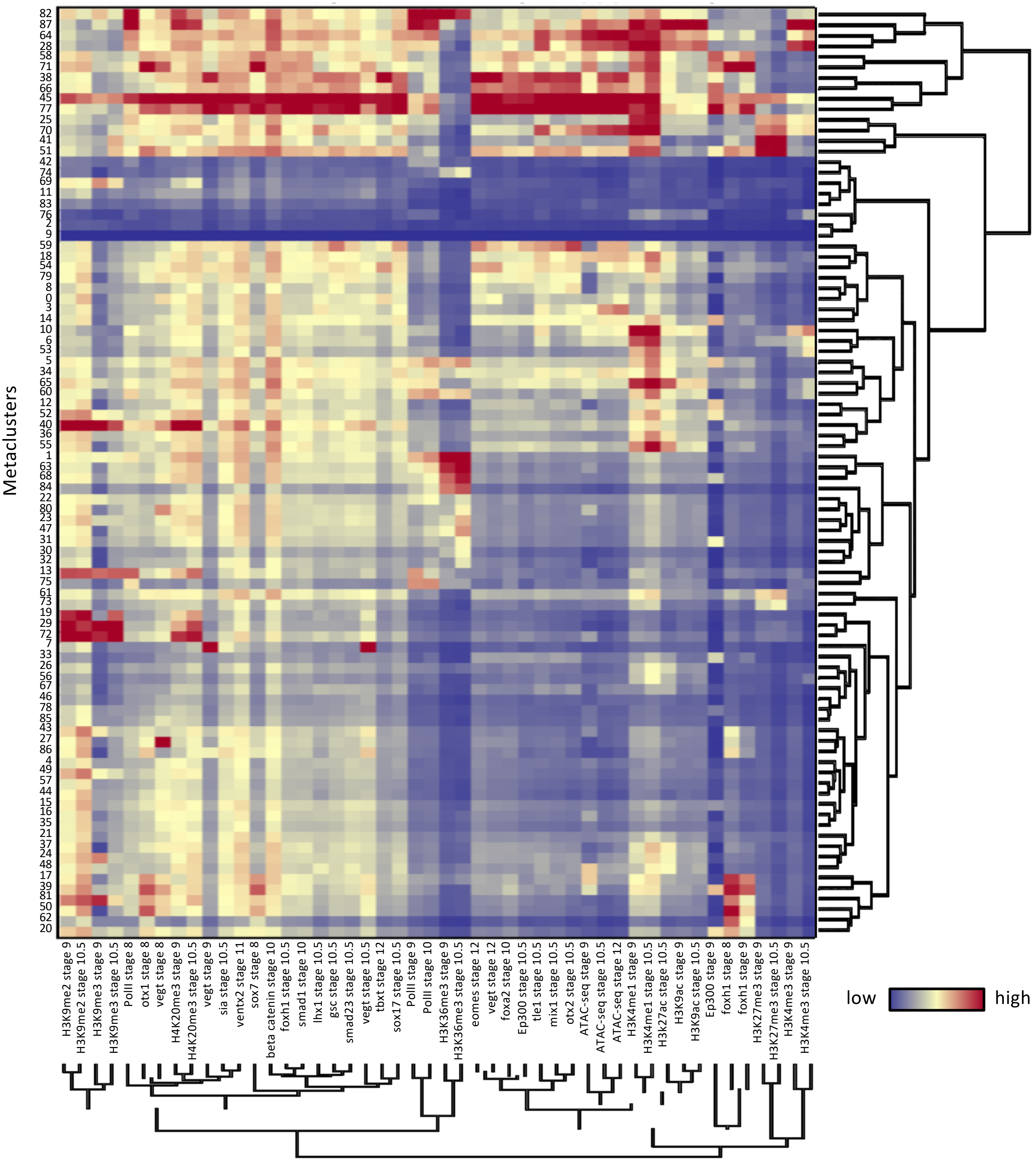


Suppl Figure 3

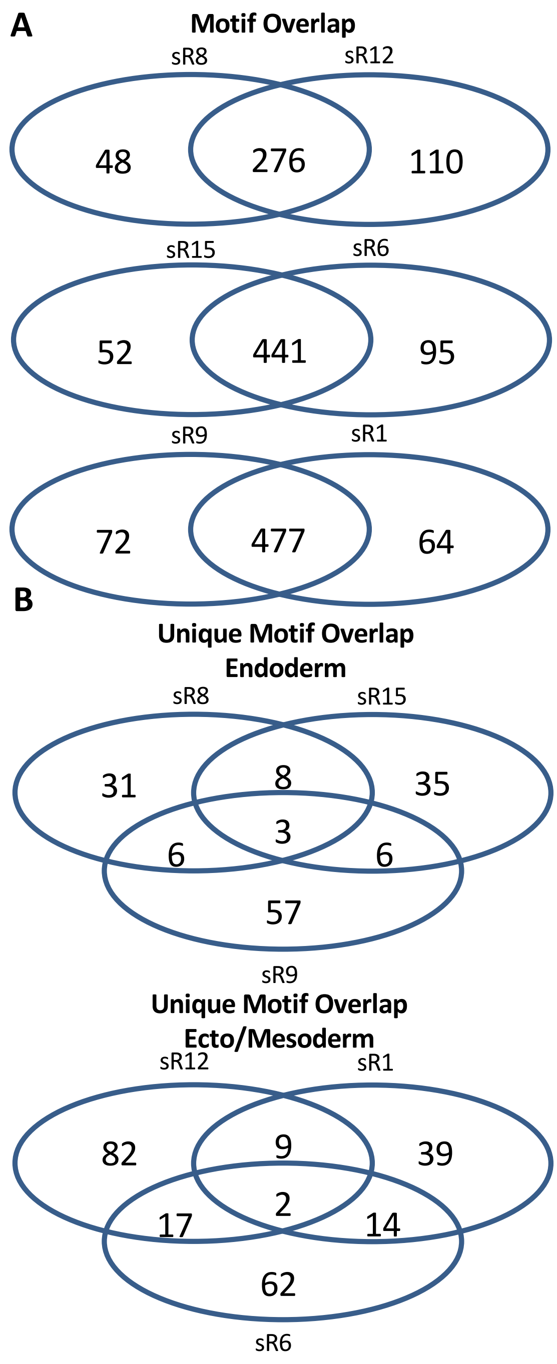


Suppl Figure 4

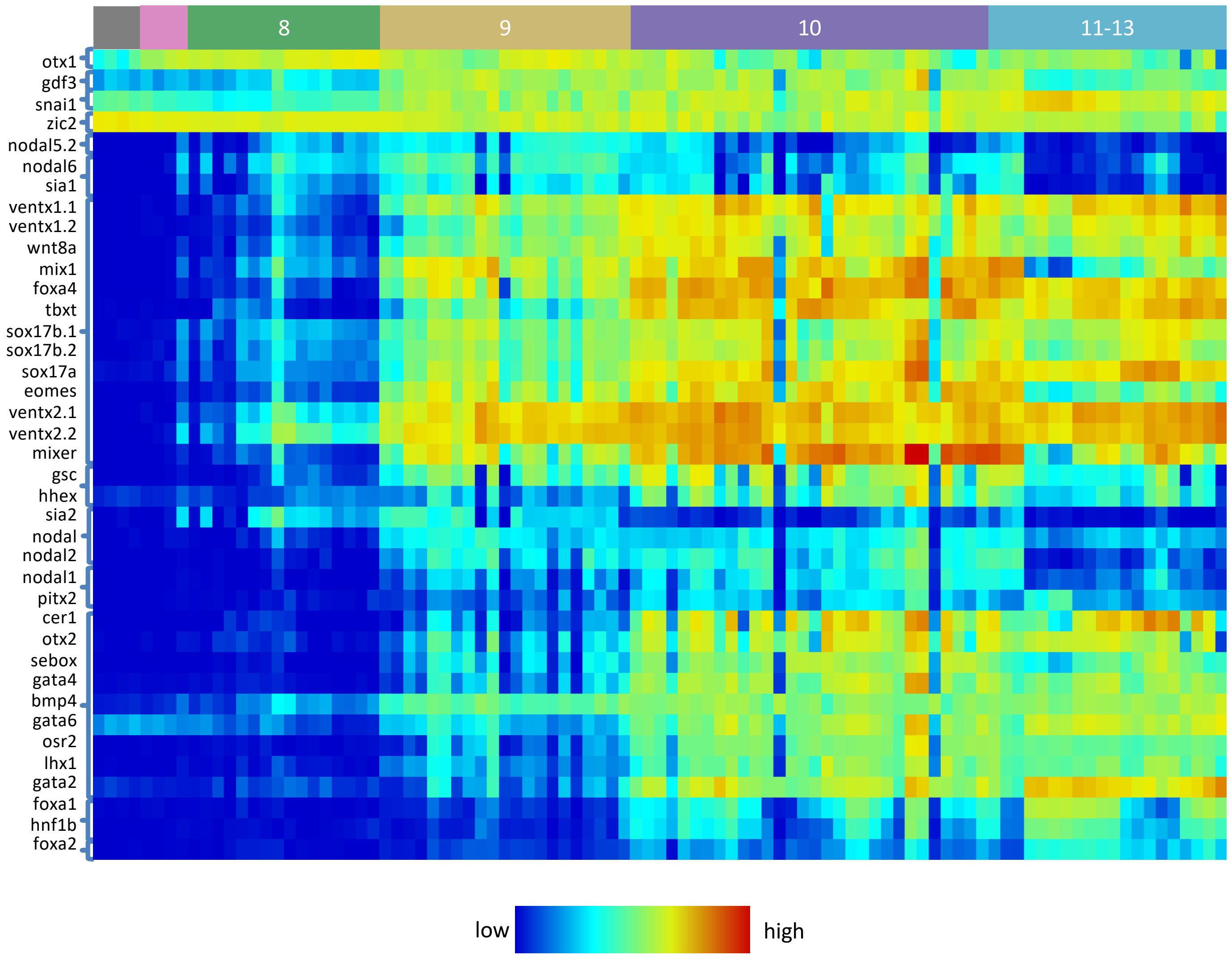


Suppl Figure 5

A

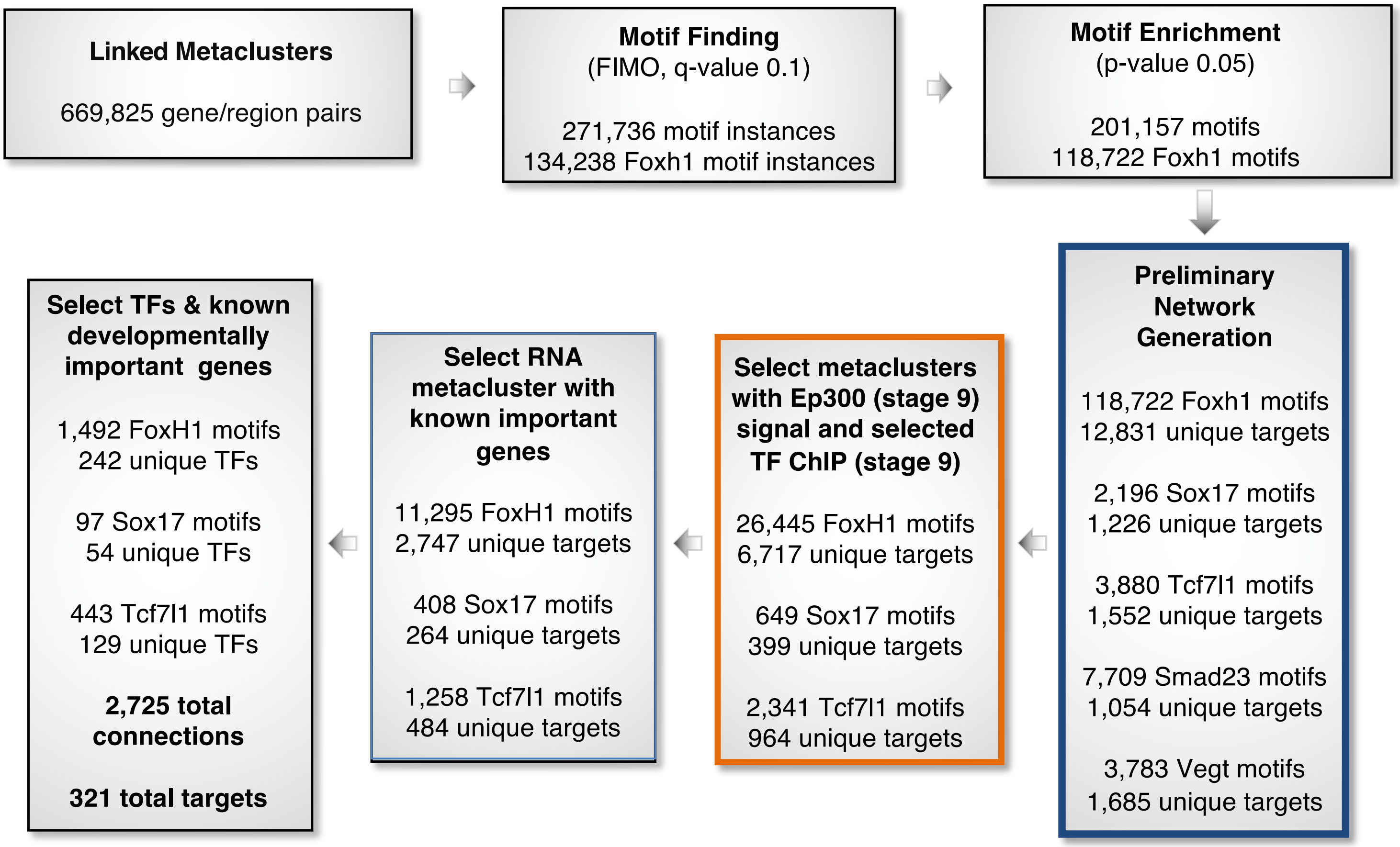


B

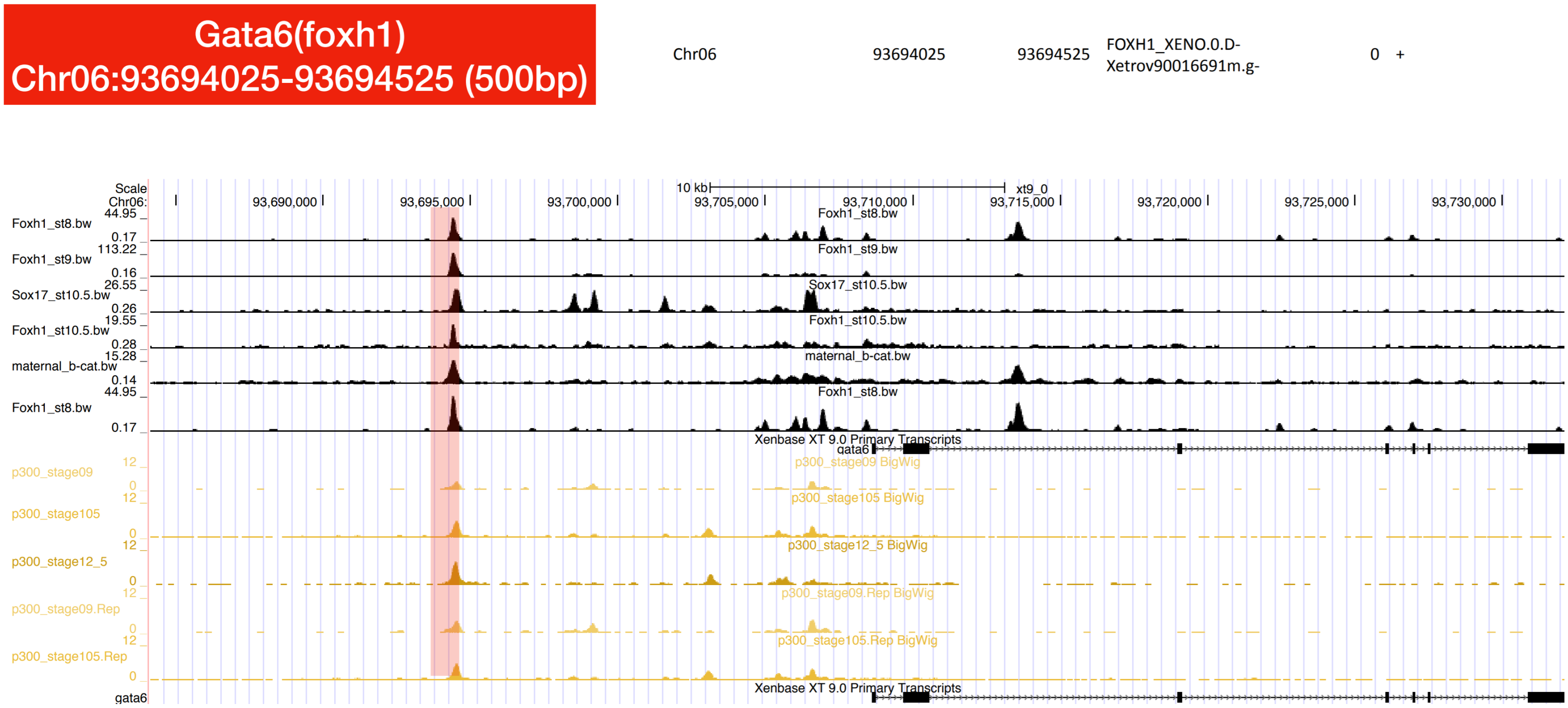


Suppl Figure 6
